# Perturbation-response analysis of *in silico* metabolic dynamics in nonlinear regime: Hard-coded responsiveness in the cofactors and network sparsity

**DOI:** 10.1101/2023.10.18.562862

**Authors:** Yusuke Himeoka, Chikara Furusawa

## Abstract

Homeostasis is a fundamental characteristic of living systems. Unlike rigidity, homeostasis necessitates that systems respond flexibly to diverse environments. Understanding the dynamics of biochemical systems when subjected to perturbations is essential for the development of a quantitative theory of homeostasis. In this study, we analyze the response of bacterial metabolism to externally imposed perturbations using kinetic models of *Escherichia coli*’s central carbon metabolism in nonlinear regimes. We found that three distinct kinetic models consistently display strong responses to perturbations; In the strong responses, minor initial discrepancies in metabolite concentrations from steady-state values amplify over time, resulting in significant deviations. This pronounced responsiveness is a characteristic feature of metabolic dynamics, especially since such strong responses are seldom seen in toy models of the metabolic network. Subsequent numerical studies show that adenyl cofactors consistently influence the responsiveness of the metabolic systems across models. Additionally, we examine the impact of network structure on metabolic dynamics, demonstrating that as the metabolic network becomes denser, the perturbation response diminishes—a trend observed commonly in the models. To confirm the significance of cofactors and network structure, we constructed a simplified metabolic network model, underscoring their importance. By identifying the structural determinants of responsiveness, our findings offer implications for bacterial physiology, the evolution of metabolic networks, and the design principles for robust artificial metabolism in synthetic biology and bioengineering.

## Introduction

For bacterial cells steadily growing under fixed, substrate-rich culture conditions, several quantitative laws pertaining to the physiology of growing bacteria have been identified. These include the linear association between growth rate and the ribosomal fraction in total proteomes Schaechter et al. (1958); Scott et al. (2010), the linear relationship linking substrate uptake to growth rate Pirt (1965), and the correlations between transcriptional responses to various sublethal stresses Kaneko et al. (2015).

In recent years, the dynamical aspects of bacterial physiology has garnered significant attention. A wealth of intriguing phenomena under such conditions has been gradually revealed. For instance, cells can “remember” the duration of starvation and the nature of environmental changes—such as the rate at which nutrients deplete—and this can dramatically alter the distribution of lag times Kaplan et al. (2021); Himeoka and Kaneko (2017); Levin-Reisman et al. (2010); Himeoka et al. (2022a). Both the proteome and metabolome of starved cells display characteristically slow dynamics Radzikowski et al. (2016). The metabolites used for storage play a pivotal role in fluctuating environments Sekar et al. (2020). Further, there is a trade-off between the growth rate under nutrient-rich conditions and other physiological parameters: namely, the lag time upon nutrient shifts Basan et al. (2020) and the death rate under starved conditions Biselli et al. (2020).

To achieve a comprehensive understanding of physiological responses, the development of quantitative theories that elucidate the dynamics of cellular physiology is indispensable. Metabolism, often a primary trigger for physiological transitions, stands at the forefront of efforts to forge a unified perspective on the dynamics of bacterial physiology. To this end, extensive research has been undertaken to understand the dynamic responses of metabolic systems using mathematical frameworks; The metabolic control analysis (MCA) Heinrich and Rapoport (1974), the biochemical systems theory (BST) especially S-systems Savageau (1988), and the linear stability analysis of massbalance kinetic models Chakrabarti et al. (2013); Lee et al. (2014). The exploration of the study on the stability and responsiveness of the metabolic system is limited only to the linear region except S-systems. However, the S-system approach omits the mass-balance. The mass-balance condition, or in other words, the stoichiometric constraints, is known to have strong impact on the dynamics of chemical reaction systems Feinberg (2019). For the development of theoretical foundations of metabolic dynamics, it is essential to explore the dynamic responses of mass-balancing metabolism beyond the linear regime.

In this manuscript, we examine the mass-balancing kinetic models of *Escherichia coli*’s central carbon metabolism and ask how the metabolic state responds to perturbations with which the linear approximation is no longer valid. We study three independently proposed kinetic models Chassagnole et al. (2002); Khodayari et al. (2014); Boecker et al. (2021), and explore the features shared among the models. We demonstrate that the three models exhibit strong responses to perturbations on the metabolic states; The effects of these perturbations are magnified, leading to significant deviations in metabolite concentrations from their steady-state values, depending on which and how metabolite concentrations are perturbed. The observed responsiveness is a hallmark of the realistic metabolic network as the toy metabolic models lack such a strong response. Computational analysis revealed that cofactors, such as ATP and ADP, play a crucial role in the strong response to perturbations. We also discovered the importance of the sparse structure of the metabolic network by adding virtual reactions to it. To validate our findings, we developed a simple, minimal model of metabolic reactions, and we confirmed that cofactor dynamics and network sparsity are indeed the key ingredients of the strong responses.

## Results

### Model

In the present study, we investigate the dynamic response of metabolic states to perturbations in metabolite concentrations using kinetic models of *E*.*coli* central carbon metabolism. In kinetic modeling approaches, the temporal evolution of metabolites’ concentrations is modeled using ordinary differential equations to capture behaviors in out-of-steady-state metabolism.

We employ three kinetic models of *E*.*coli*’s central carbon metabolism, emphasizing the common features exhibited by the models. A limitation of kinetic models is their specificity; unlike constraint-based modeling, kinetic modeling necessitates extensive biochemical information, such as the function form of the reaction rate equation and the parameter values for each reaction. By examining a single kinetic model, conclusions may apply only to that specific model. Therefore, our focus lies on the shared features of the three models to circumvent the limitations of individual model specificity. We also believe there is potential for the results obtained commonly among the models to apply to real biological systems as well.

The models under consideration are those proposed by Chassagnole et al. Chassagnole et al. (2002), Khodayari et al. Khodayari et al. (2014), and Boecker et al. Boecker et al. (2021). While all three models incorporate the glycolytic pathway, only the models by Khodayari et. al. and Chassagnole et. al. feature the pentose phosphate (PP) pathway. Notably, the Chassagnole model excludes the tricarboxylic acid (TCA) cycle. A graphical summary of the metabolic modules in these models is presented in **Fig. 1A**.

**Figure 1.**
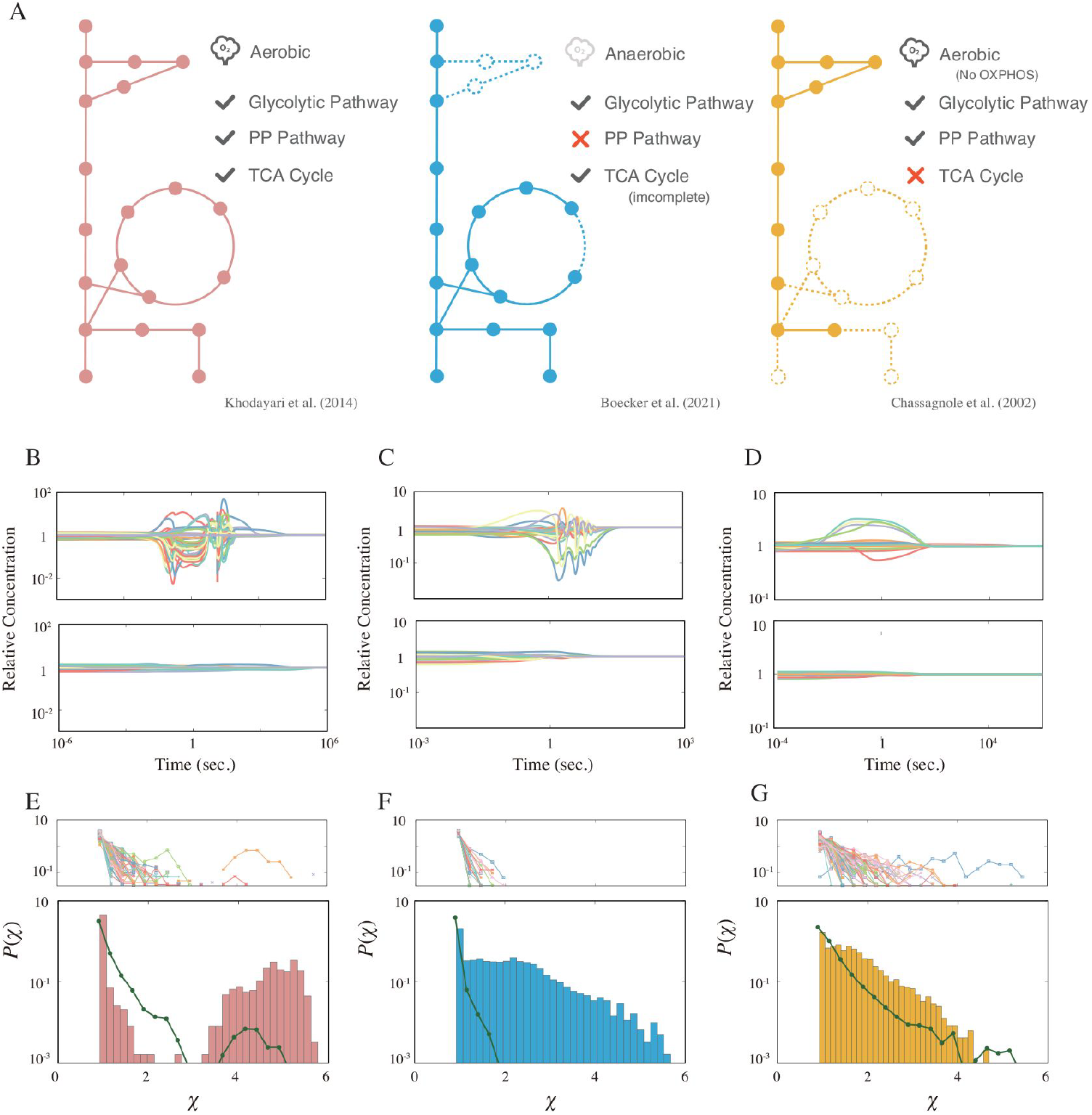
(A) A graphical summary of the three models used in this study. The solid circles represent the metabolites and the lines represent the metabolic reactions. The dashed circles and lines are the subsystems not implemented in the model. (B-D). Example time courses for each model. The top and bottom panels are examples of the strong and weak response, respectively. (E-G). The response coefficient distribution *P* (*χ*) of each model (bottom). The average response coefficient distribution of the random catalytic reaction network (RCRN) model is overlaid (green line). The response coefficient distribution of each instance of the RCRN model is shown on the top panel. The model schemes, time courses, and distributions are aligned in each column, i.e. the leftmost network of (A), (B), and (E) are from the model developed by Khodayari et al. Khodayari et al. (2014).

The Boecker and Chassagnole models utilize glucose as the sole carbon source, whereas the Khodayari model can assimilate other carbon sources, such as fructose, formate, and acetate depending on differences between the extracellular- and intracellular concentrations. The Boecker model explicitly models biomass formation, considering several metabolites as biomass precursors that are consumed with a fixed stoichiometry to produce biomass. Conversely, the Khodayari and Chassagnole models do not incorporate the biomass formation reaction. Instead, biomass precursor metabolites are consumed independently, implying that the stoichiometry of biomass formation varies depending on the concentrations. The dilution effect due to the volume growth is modeled in both the Boecker and Chassagnole models. In the Boecker model, the dilution rate corresponds proportionally to the biomass formation reaction rate, while it remains constant in the Chassagnole model. The Khodayari model omits the dilution effect. In addition to the metabolic reaction implemented in each model, the specific forms of the reaction rate equations also differ among the models.

Within these models, the dynamics of gene regulation mediated by transcription factors are not modeled; that is, the total concentration of each metabolic enzyme is constant while the substralevel regulations are incorporated into the models (The lists of substrate-level regulations are provided in **Table. S1–S3**. For a detailed description, see *Materials and Methods*).

### Perturbation-response simulation

In this section, we present the results of a series of simulations, which we call perturbation-response simulations. These simulations highlight the unique responses of the metabolic models to perturbations and shed light on the dynamic behavior of metabolic systems. It may be useful for the reader to first describe the perturbation-response simulation procedure. The procedure is consistently applied across all three models used in our study and can be summarized as follows:

1. Compute the growing steady-state attractor.
2. Generate *N*_ini_ initial points by perturbing the metabolite concentrations from the attractor.
3. Simulate the model dynamics starting from each initial point.

#### 1. Computing the attractor

First, we identify the steady-state attractor where the production/consumption of each metabolite is balanced. For the Khodayari and Boecker models, the attractor corresponds to the steady-state studied in their original papers. For the Chassagnole model, we numerically determined the steadystate attractor (details provided in *Materials and Methods*).

#### 2. Generating the initial points by perturbation

Once the steady-state attractor has been computed, we establish a set of initial points for subsequent computations by perturbing the metabolite concentrations. This perturbation is a proxy for stochastic fluctuations within cells. We postulate that the perturbation source in the variability in protein concentrations due to the inherent randomness of transcription and translation processes. Also, the cell division would be another source of perturbations of metabolites’ concentrations while the cell division is not explicitly considered in the study. Based on a stochastic model of transcription and translation with a biologically relevant setup, the relative size of the concentration fluctuations is estimated to be several tens of percent (for a detailed derivation, see *Materials and Methods*). Consequently, the perturbed concentration of chemical *n* as 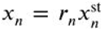, where *r*_*n*_ is a uniformly distributed random number with a maximum strength of 40%, i.e. ranging from 0.6 to 1.4. The perturbation strength is beyond the linear region that the relaxation dynamics is approximated by the linearlized model around the steady state. The choice of uniform distribution is not essential for the following result. Also, the characteristic features of the response studied later remain consistent even when different perturbation strengths are employed (see **Fig. S2**).

#### 3. Simulating the dynamics

We compute the model dynamics using the *N*_ini_ points generated by the perturbation as initial points to explore the metabolic response to the perturbations. As far as we have tried, all initial points have returned to the original growing steady state, and thus, we hereafter study the relaxation behavior.

### Strong responses in the kinetic models

After executing the perturbation-response simulations for the models, we observed distinct relaxation dynamics in each. In **Fig. 1B-D**, the top and bottom panels display typical examples of dynamics from each model that either respond strongly or weakly to perturbations. In the weakly responding dynamics (bottom panel), the effects of the perturbations diminish almost monotonically. Conversely, in the dynamics that respond strongly (top panel), the initial displacements of concentrations are amplified over time, leading some metabolites’ concentrations to overshoot or undershoot (significant concentration changes in metabolites are highlighted in **Fig.S3**).

For more in-depth analysis, we introduce a measure of the responsiveness to the perturbation that we call the response coefficient *χ*, defined as

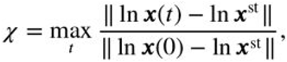

where ‖⋅‖ is the Euclidean norm. The response coefficient distributions for each model are plotted in the bottom panels of **Fig. 1E-G**. All three models have the peak at *χ* = 1 (note that this is the minimum value of *χ*). These peaks correspond to the trajectories where the Euclidean distance between the state and the attractor decreases monotonically with time (i.e. max_*t*_ ‖ ln ***x***(*t*) − ln ***x***^st^ ‖ = ‖ ln ***x***(0) − ln ***x***^st^ ‖). In addition, the distribution of the Khodayari model exhibits another peak in a larger *χ* region while the other two models have plateaus adjacent to the primary peak. We have computed the response coefficient by linearizing each model. As shown in **Fig. S1**, the responsive-ness of the original model cannot be described by the linearized model.

For comparison purposes, we conducted perturbation-response simulations for the random catalytic reaction networks (RCRN) model, a toy model of the metabolic network Furusawa and Kaneko (2003, 2012). In one instance of the RCRN model, the “metabolites” are interconnected through a random network, and every metabolite is considered a catalyst as well as a reactant. We generated 128 instances of the RCRN model, each having the same number of metabolites and the same distribution of the reaction rate constant. For instance, to draw a comparison with the Khodayari model, we generated the model instances with 778 metabolites and 3112 reactions, where the reaction rate constants are randomly chosen from that of the Khodayari model (detailed construction of the RCRN model is provided in *Materials and Methods*). The top panel of **Fig. 1E-G** presents the response coefficient distribution for each RCRN model instance, represented by different lines. Additionally, the “average” response coefficient distribution, which is the mean of the distributions plotted in the top panel, is overlaid (green dotted line). As a fact, the emergence of the additional peak and plateaus, indicative of strong responsiveness, are unique traits of the realistic metabolic reaction network model and are seldom seen in toy representations of the metabolic network.

### Key metabolites on the responsiveness

What factors contribute to the strong responses of the metabolic state? In this section, we investigate which metabolites play pivotal roles in inducing these strong responses in the model. To understand the role of each metabolite’s dynamics, we conducted perturbation-response simulations using model equations derived by fixing each metabolite’s concentration to its steady-state value, i.e, set 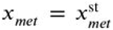 and *dx*_*met*_/*dt* = 0 for the concentration of the metabolite *met* whose con-centration is fixed. The impact of holding each metabolite’s concentration constant is quantified by the change in the average response coefficient:

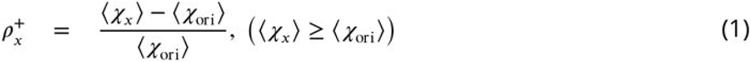

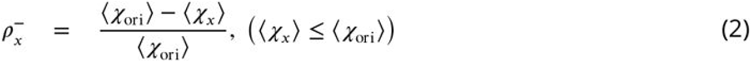

where *x* denotes the metabolites’ IDs and *χ*_*x*_ represents the response coefficient of the model with the concentration of the metabolite *x* fixed to constant. Note that for each metabolite in each model, either the 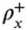 or 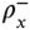 is defined, depending on whether the modified model (with the metabolite *x* as a held constant) exhibits an average response coefficient larger (for 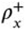) or smaller (for 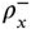) than the original model.

**Fig. 2** displays 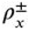 values for representative metabolites (comprehensive results for all metabolites can be found in **Fig. S4**). While most metabolites have a bilateral effect on the average response coefficient depending on the original model (or exert a small unilateral effect), ATP and ADP (we refer to them collectively as AXPs) consistently demonstrate a marked impact in reducing the average response coefficient across models. This observed significance of AXPs in metabolic dynamics aligns with previous findings Himeoka and Mitarai (2022). The common importance of AXPs are shown also when we quantify the impact of their dynamics on the responsiveness by utilizing the Sobol’ total sensitivity index (**Fig. S5**) Homma and Saltelli (1996).

**Figure 2.**
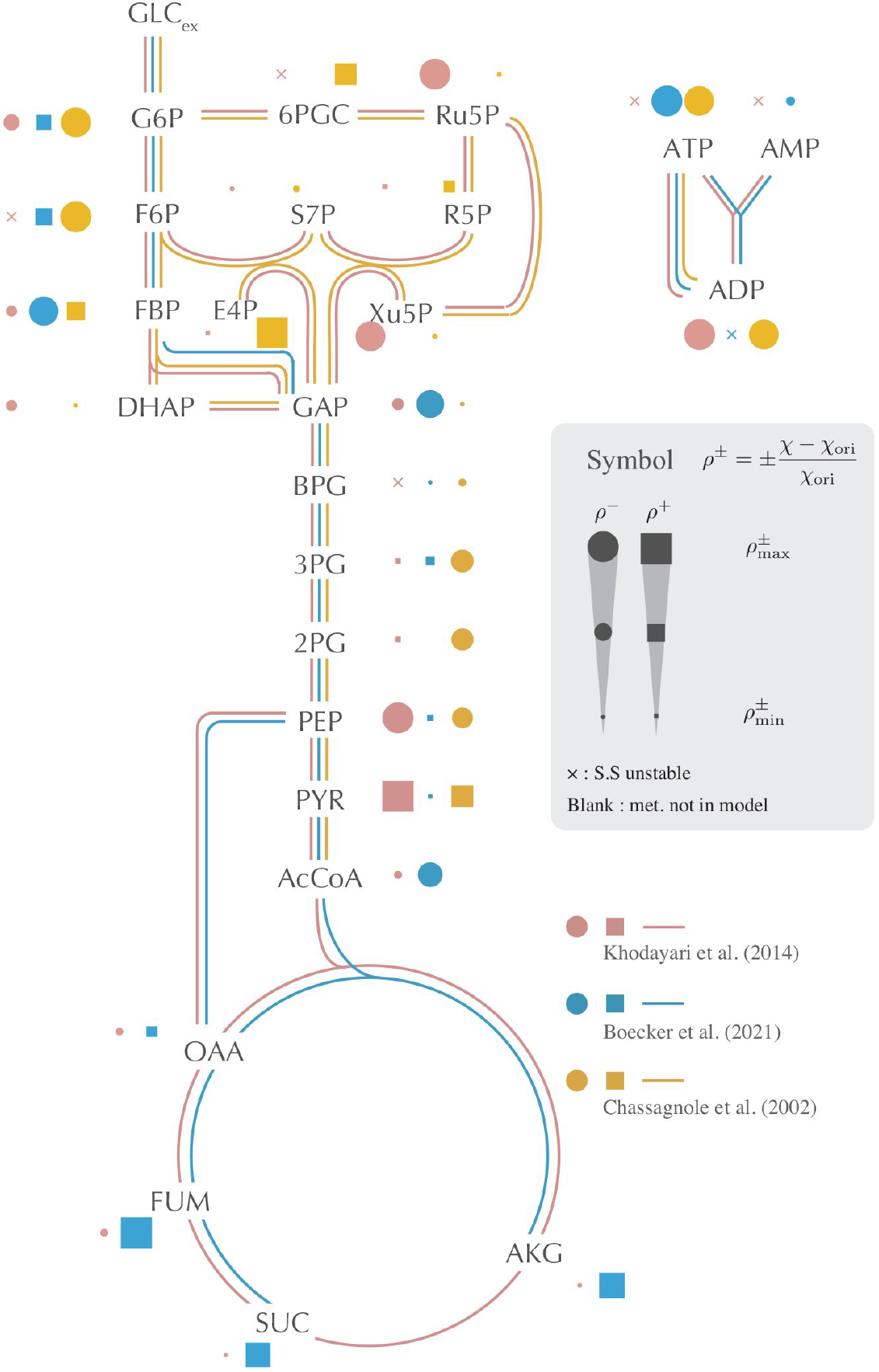
The impact of setting each metabolite’s concentration to its constant, steady-state value is depicted on the metabolic network. This impact is quantified by the relative change in the average response coefficient, as detailed in (1) and (2). The filled point and filled square symbols represent the decrease 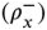 and the increase 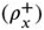 of the response coefficient, respectively. The size of the symbols is scaled using the maximum and minimum values of 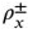 computed for each model. The cross symbol indicates that the original steady-state attractor becomes unstable when the concentration of the corresponding metabolite is fixed. If a dot, square, or cross is absent, the concentration of the corresponding metabolite is not a variable in the model. Only selected, representative metabolites are displayed in the figure. Comprehensive results are provided in **Fig. S4**.

Interestingly, halting the dynamics of several metabolites can amplify the model’s response coefficient. Across the models, metabolites that augment the response coefficient when their concentrations are fixed tend to be allosteric regulators associated with negative regulation. Freezing these metabolites’ concentrations effectively removes the negative feedback loop present in the metabolic systems. Typically, negative feedback stabilizes system behavior. In the present context, it assists the system in returning to its original state after perturbations. Hence, fixing these metabolites’ concentrations allows the system to respond more vigorously to perturbations compared to the original model. We checked the effect of feedback itself on the responsiveness rather than the concentration dynamics by weakening the strength of substrate-level regulation of OAA, FUM, AKG in the Boecker model (**Fig. S6**). It was shown that weakening the substrate-level regulation of CSICD by AKG and MDH by OAA increases the responsiveness. On the other hand, weakening the strength of substrate-level regulation of neither FRD by OAA nor MDH by FUM alters the responsiveness, and in addition there is no substrate-level regulation by SUC. The increase in responsiveness by fixing the concentrations of OAA and AKG would be explained by substrate-level regulation of the corresponding metabolites. However, substrate-level regulation is not sufficient to describe the effect of fixing the concentration of FUM and SUC. Higher order effects may be required.

Pyruvate is an exception to the above description, which is not the regulator in the models. The strong influence of pyruvate on the increase in response coefficients is attributed to the phosphotransferase system (PTS). The external glucose is taken up by converting phosphoenolpyruvate into pyruvate through the PTS. This reaction elevates pyruvate concentration, subsequently decelerating glucose uptake. By holding pyruvate concentration constant, this negative feedback effect is negated, enhancing the autocatalytic nature of the reaction system formed by the PTS and glycolysis (GLC_ex_ + PEP → … → 2PEP).

### The role of the sparsity of the networks

The key role of AXPs in changing the kinetic models’ responsiveness was shown in the previous section. Nevertheless, this does not imply that all kinetic models of chemical reaction systems with cofactors such as AXPs always exhibit a strong response to perturbations. The strong responses demonstrated by the three models (as seen in **Fig. 1B-D**) are absent in abstract toy models of metabolism, which consist of random reaction networks incorporating such cofactor chemicals Kondo and Kaneko (2011). This discrepancy prompts us to investigate other potential determinants causing metabolic models to display strong responses shown in the three models.

A noteworthy attribute of metabolic networks is their inherent sparsity. While cofactors are linked to numerous reactions, the backbone networks—networks excluding cofactors—are distinctly sparse. As an example, the glycolytic pathway predominantly mirrors a linear sequence of reactions, resulting in many metabolites within this pathway participating in merely two reactions.

Does network structure play a role in the dynamics of metabolic systems? For this question, previous research in the opposite direction—reaction dynamics on dense networks—provides useful insights. There are several studies using the random networks as an abstract model of metabolic networks Furusawa and Kaneko (2003); Kaneko et al. (2015); Awazu and Kaneko (2009); Himeoka et al. (2022b). In such types of models, the effect of the initial perturbations on metabolite concentrations typically decays monotonically over time Awazu and Kaneko (2009); Himeoka et al. (2022b). This is in stark contrast to the behavior observed in **Fig. 1B-D**.

Considering the inherent tendency of random reaction networks to display dynamics characterized by weak responses, we are compelled to investigate the influence of network sparseness on the dynamic behavior of metabolic models. To this end, we increase the reaction density of the metabolic network in the three models and assess the resultant changes in responsiveness. Our approach entails initially introducing *N*_add_ random reactions into the network and subsequently al-locating kinetic parameters to these additions from the distribution of parameter values computed from the original model. We then execute a perturbation-response simulation on this extended model to determine the response coefficient, which in turn helps ascertain the distribution and average of these coefficients. This process is schematically depicted in **Fig. 3A**.

**Figure 3.**
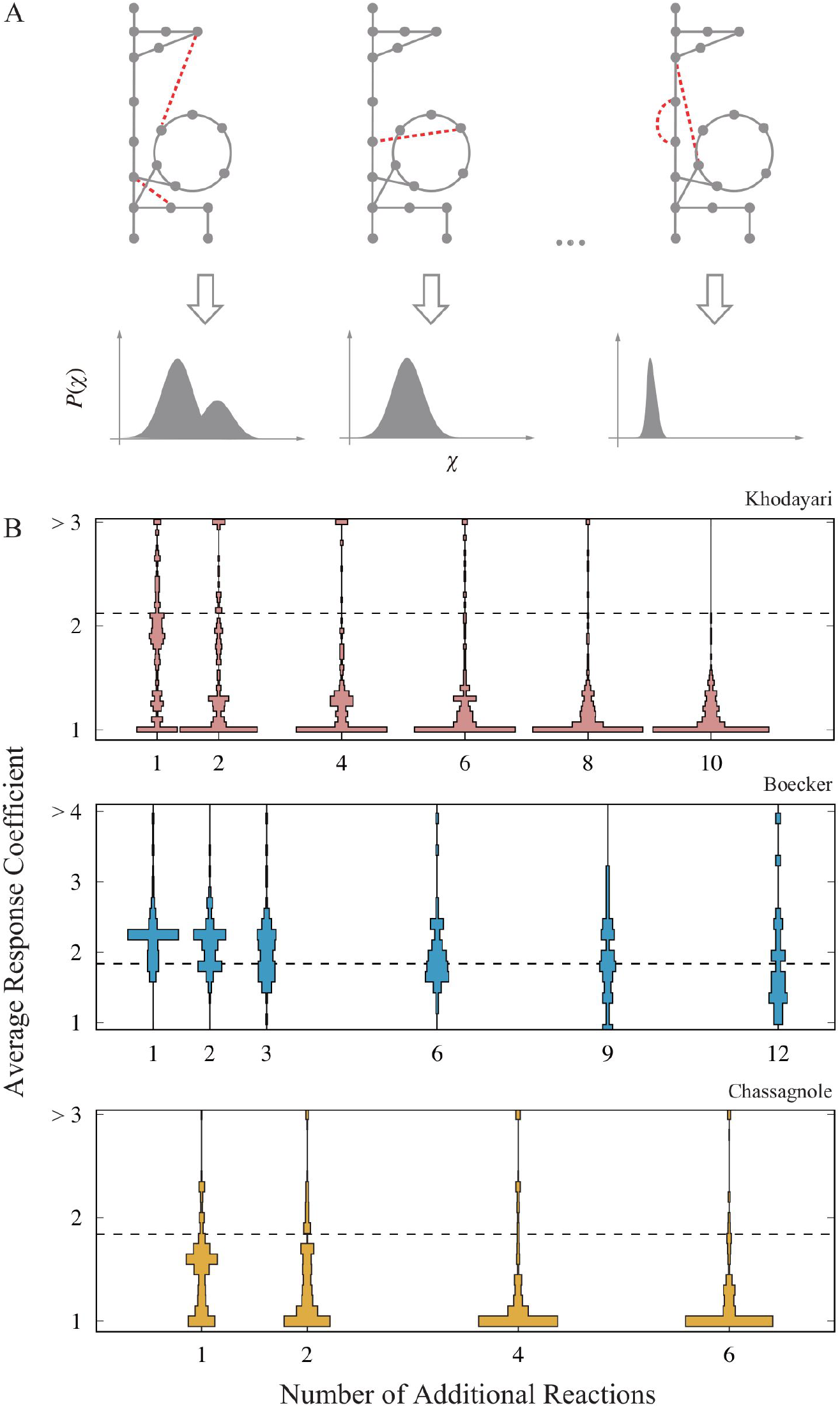
(A) schematic figure of the network expansion simulation. Reactions are randomly added to the original metabolic network, and then, the corresponding ODE model is constructed. In the illustration, the additional reaction is highlighted as the dashed red lines. The perturbation response simulation is performed on the expanded models to obtain the response coefficient distribution. The average response coefficient is calculated as a scalar indicator of the responsiveness for each response coefficient distribution. (B). The distribution of the average response coefficient is plotted against the number of additional reactions for the three models. The dashed line indicates the average response coefficient of the original model with no additional reaction. For each *N*_add_, we generated 256 extended network, and 128 trajectories were computed for each extended network.

**Fig. 3B** illustrates the alterations in responsiveness resulting from the model extensions. We generated extended metabolic networks and accompanying ODE models by adding *N*_add_ random reactions, and executed the perturbation-response simulation. The average response coefficient is then computed for each model and the distribution of the average response coefficient is presented as the violin plots. In the perturbation-response simulation, the extended models with only a single attractor are subjected to the response coefficient analysis (comprehensive procedural details available in the *Materials and Methods* section). Note that in this procedure, metabolites are not newly introduced to the model, but only the reactions are. Thus, the reaction density increases.

As evident from the figures, a consistent trend emerges across the three models: the introduction of random reactions weakens responsiveness. However, the magnitude of this reduction varies among the models. It is important to note that the reactants of the added reactions are selected randomly, thus not based on any specific chemical rationale. Nevertheless, this diminished responsiveness persists even when the additional reactions are confined solely to those cataloged in the metabolic model database, as visualized in **Fig. S7**.

Such attenuation aligns with expectations shaped by prior studies on dense reaction networks Furusawa and Kaneko (2003); Kaneko et al. (2015); Awazu and Kaneko (2009); Himeoka et al. (2022b). Evidently, the inherent sparsity of the backbone reaction network plays a critical role, enabling the kinetic model of metabolic systems to respond strongly to perturbations.

### A minimal model for disrupted homeostasis

So far, we have studied the kinetic models of the central carbon metabolism of *E*.*coli*. Our findings indicate that cofactors and the sparsity of the backbone network (network devoid of cofactors) are the keys to the responsiveness of the metabolic dynamics. In this section, we develop a simple minimal model to investigate if these two factors are sufficient to induce strong responses in reaction systems.

Our minimal model is designed in a manner that allows us to adjust the sparsity of the backbone network and the proportion of cofactor-coupled reactions. First, we explain the construction of the backbone networks. These networks are comprised of *N* chemical species, C_1_, C_2_, …, C_*N*_, and *R* reactions (*R* ≥ *N* −1). For simplicity, we construct the backbone network using only uni-uni reactions. The minimum number of uni-uni reactions required to interlink *N* chemical species is *N* − 1. To assemble a backbone network with *N* chemicals and *R* reactions, we first connect *N* reactions using *N* − 1 reactions and subsequently introduce random reactions for the pairs of chemicals that lack reactions, totaling *R*−(*N* − 1). We incorporate exchange reactions with extracellular environments for 5% of the metabolites (this percentage is not critical to the results). The exchange reactions are always introduced to the two endpoints metabolites of the sequential linkage to ensure there are no dead-ends on reaction networks. We posit that one of the endpoint metabolites from the sequential linkage is the nutrient metabolite, and we set its external concentration at 100, while for the remainder, it is set to unity.

As cofactors, we introduce the three forms of the activation for mimicking AXP, A^**^, A^*^ and A, which are not included in the *N* chemicals in the backbone network. We introduce the coupling fraction *f* to modulate the number of reactions with cofactor coupling. With the coupling fraction *f*, the number of reactions with cofactor coupling is given as Round(*fR*) with Round(⋅) as the function to round the value to the nearest integer. When a reaction is selected for cofactor coupling, two forms of the cofactor (chosen from A^**^, A^*^ and A) are randomly selected. The reaction then gets coupled to the conversion between these two cofactor forms, with the direction being randomly set.

Overall, the differential equation governing the minimal model can be expressed as

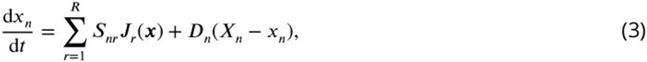

In this expression, the concentrations of the chemicals from the backbone network, as well as the cofactors, are combined in a vertical stack, i.e., *n* = 1, 2, …, *N, N* + 1, *N* + 2, *N* + 3. *x*_*n*_ with *n* = 1, …, *N* represents the concentrations of the *n*th chemical in the backbone network, and *x*_*N*+1_, *x*_*N*+2_ and *x*_*N*+3_ denote the concentrations of A^**^, A^*^ and A, respectively. *S* is the stoichiometric matrix of the reaction network. *D*_*n*_ is unity if the *n*th chemical has an influx/efflux term, otherwise, it is zero. *X*_*n*_ is the external concentration of chemical *n*, which is a constant parameter.

Mass action kinetics is employed for the reaction rate function *J*_*r*_. The rate function has the form

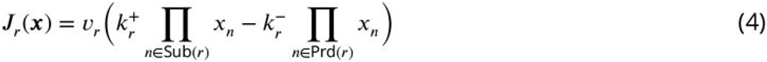

where Sub(*r*) and Prd(*r*) are the set of substrates and products, respectively, of the *r*th reaction. If the reaction is coupled with the cofactor, |Sub(*r*) | = |Prd(*r*) | = 2 and otherwise |Sub(*r*) | = |Prd(*r*) | = 1. We neither allowed the self-loops in the backbone network (C_*n*_ ⇌ C_*n*_) nor the catalytic activity of the cofactors (e.g., C_*n*_ + A^*^ ⇌ C_*m*_ + A^*^).

The parameters *v*_*r*_ and are 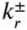 the reaction rate and irreversibility, respectively. The *v*_*r*_ values are randomly set as 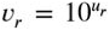 with the uniformly distributed random number, *u*_*r*_ ~ *U*(−3.66, 7.13). The range for the kinetic parameter is derived from the Khodayari model. 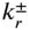 is calculated using the Arrhenius equation based on the standard chemical potential of each chemical. We randomly set the standard chemical potential *μ*_*n*_ for each chemical, where *μ*_*n*_ follows the uniform distribution *U*(0, 1), while the standard chemical potentials of A^**^, A^*^ and A are fixed to the constant values, 1.0, 0.5, and 0.0, respectively. To satisfy the detailed balance condition, 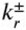 are then given by 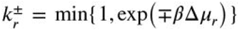 with Δ*μ* as the standard chemical potential difference between the products and the substrates, Δ*μ*_*r*_ = ∑_*n*∈Prd(r)_ *μ*_*n*_ − ∑_*n*∈Sub(*r*)_ *μ*_*n*_, and *β* is the inverse temperature.

We generate *M* networks for several choices of reaction number *R* and coupling fraction *f*. We performed the perturbation-response simulation to determine the response coefficient distribution of each network. In **Fig. 4**, the average response coefficient, averaged over the networks, is depicted as functions of *R* and *f* (see the rectangles labeled as “Cofactor”). These figures clearly illustrate that the average responsiveness tends to decrease as either *R* increases or *f* decreases. The coupling with the cofactors leads to the two-body reactions, and thus, increases the nonlinearity of the model equation. It is noteworthy that the increase in responsiveness with *f* cannot be fully attributed to the increase in the model’s nonlinearity. To validate this, we employed the model (3) without the cofactors, but incorporated catalytic reactions. In this model, each reaction proceeds with the catalytic ability of an enzyme, represented as C_*n*_ +C_*l*_ ⇌ C_*m*_ +C_*l*_ (here, autocatalytic reactions are permissible, meaning *l* can be either *n* or *m*).

**Figure 4.**
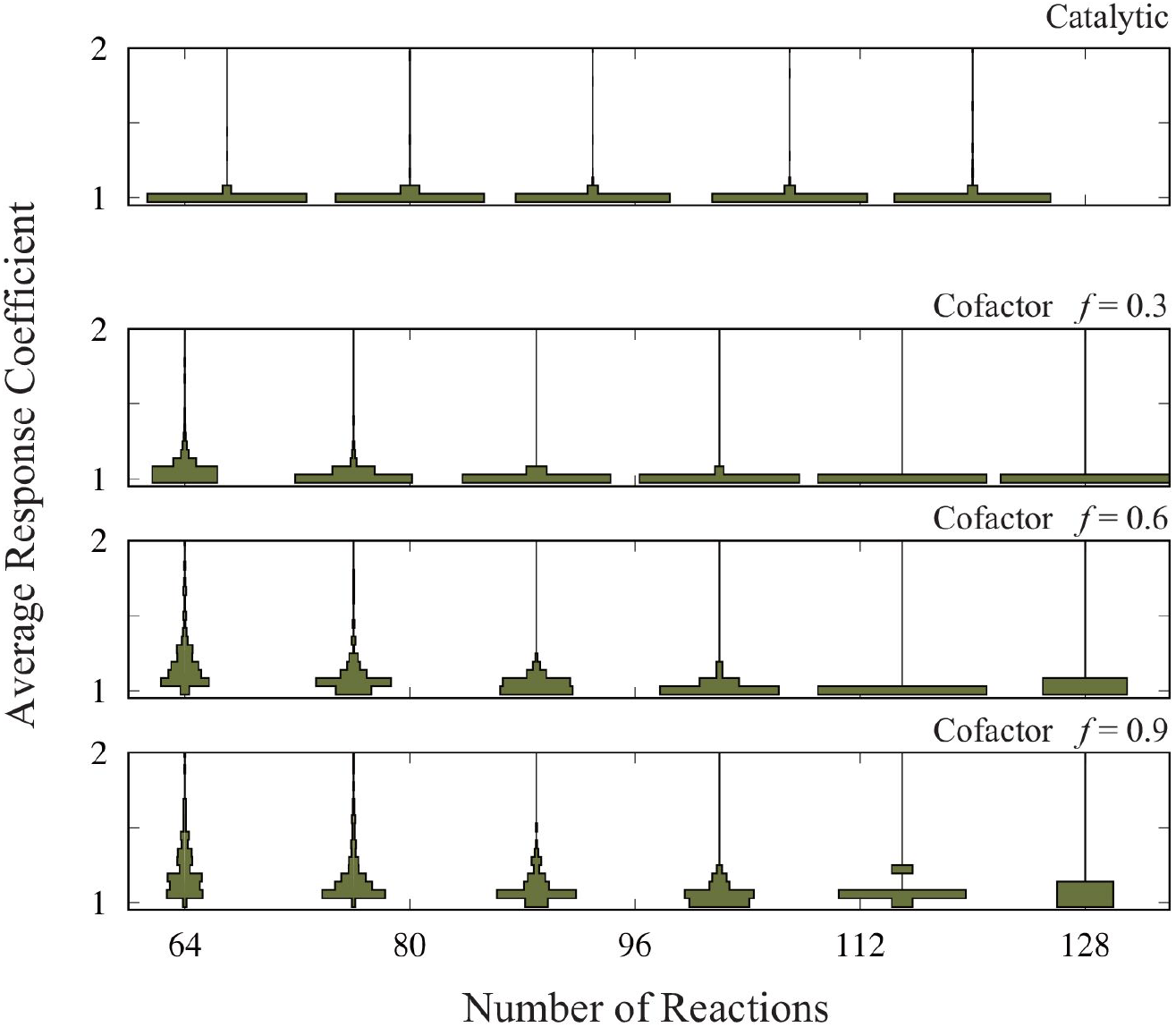
The distribution of the average response coefficient is plotted against the number of reactions, *R*, for the model with *N* = 64 metabolites plus 3 cofactors (the panels labeled as “Cofactor”) and *N* = 67 metabolites without cofactors (the panel labeled as “Catalytic”). For each combination of *R* and *f*, we generated 512 random network, and computed 128 trajectories for each of these networks. The inverse temperature *β* is set at 10 for the cofactor version and 20 for the catalytic version. This is because the maximum chemical potential difference in the cofactor version is 2.0, but it is 1.0 in the catalytic version. The total concentrations of the cofactors are set to be unity.

In terms of the numbers of constant, linear, and quadratic terms in the model equation, the catalytic reaction version of the model corresponds to the *f* = 1 case of the original minimal model. If the increase in responsiveness with *f* is dictated solely by nonlinearity, the catalytic reaction model should have comparable responsiveness with the “cofactor” model with high *f*. Yet, as shown in the rectangles above the main plot in **Fig. 4** (labeled as “Catalytic”), the responsiveness metrics are notably lower than those observed in the minimal model.

## Discussion

In the present manuscript, we have studied three models of *E. coli*’s central carbon metabolism. Our findings indicate that these models commonly exhibit strong and weak responses to perturbations of the steady-state concentration. In the weak response, the effect of the perturbation dwindles almost monotonically over time. In contrast, during a strong response, initial deviations from the steady state get amplified. Note that the linearized models cannot capture the response dynamics. We introduced the response coefficient, *χ*, as a metric to quantify the strength of these responses and undertook comprehensive computational analyses. The strong responsiveness were observed not only in the three models studied in the manuscript, but also in a model of glycolytic pathway with multiple sets of parameter choices (**Fig. S8**) Haiman et al. (2021).

We aim to identify key metabolites whose dynamics play a crucial role in the emergence of strong response. We systematically fixed the concentration of each metabolite to its steady-state value and assessed the influence of individual metabolite dynamics on the overall response strength. For a majority of metabolites, the impact on the response coefficient is bidirectional: halting the dynamic concentration of a metabolite can either amplify or diminish the response, depending upon the base model. Even if their effects consistently amplify or diminish the response, the magnitude of change in the average response coefficients remains relatively modest.

However, three metabolites—ATP, ADP, and pyruvate—stood out. They consistently and significantly influenced the average response coefficient across models. Pyruvate’s pronounced impact can be attributed to the phosphotransferase system, whereas the effects of ATP and ADP seem to arise synergistically across multiple reactions.

One of the authors has recently reported a kinetic model of *E*.*coli* metabolism to exhibit anomalous relaxations and the role of AXPs in the relaxations Himeoka and Mitarai (2022). In the paper, they reduced the metabolic network into a coarse-grained minimal network. The minimal network revealed that, depending on the initial conditions, a futile cycle conversing ATP into AMP be overly active. Then, the futile cycle can deplete ATP and ADP, leading to a significant slowdown or even cessation of several ATP/ADP-coupled reactions, pushing the entire metabolic network into a “choked” state. The extensive connectivity of AXPs is a common trait in metabolic models. Hence, it is plausible that the impact of AXPs’ dynamics on the entire metabolic system is a common feature irrespective of specific kinetic models.

Several experimental studies underscore the pivotal role of AXPs in metabolic homeostasis. For example, Teusink et al. demonstrated that in yeast, a sudden increase in glucose levels in the culture medium can activate a positive feedback loop in ATP production in the glycolytic pathway, leading to growth arrest van Heerden et al. (2014); Teusink et al. (1998). This phenomenon of growth arrest, triggered by a nutrient level surge, has been observed across various bacterial species, including *E*.*coli* Strange and Hunter (1966); Postgate and Hunter (1964); Calcott and Postgate (1972), though its molecular mechanism has been most extensively studied in *S. cerevisiae*. Moreover, a rapid decrease in the total adenyl cofactor concentration as a cellular response to external glucose up-shift has been documented Theobald et al. (1997); Kresnowati et al. (2006); Rizzi et al. (1997); Walther et al. (2010). Walther et al. found that inhibiting this transient reduction in the adenyl cofactor pool increased the number of cells transitioning to a growth-arrested state upon glucose up-shift Walther et al. (2010).

Next, we focused on the structure of the backbone network as suggested by the relatively simple dynamics typically observed in the random network metabolic models Furusawa and Kaneko (2003); Kaneko et al. (2015); Awazu and Kaneko (2009); Himeoka et al. (2022b). To this end, we increased the density of the backbone network in the three models by randomly selecting metabolites and introducing hypothetical reactions between them. The perturbation-response simulation on the extended models indicated that as more reactions were integrated, the system’s response weakened.

Finally, we devised a minimal model to evaluate our hypothesis that the dynamics of cofactors and the sparsity of the backbone network are pivotal determinants of responsiveness. This model successfully showed a strong response to perturbations when the fraction of reactions to be coupled with cofactors becomes high and the backbone network is sparse. It is worth noting that the heightened response due to cofactor coupling is not solely a result of the model equation’s increased nonlinearity. Interestingly, the model’s catalytic reaction variant demonstrated attenuated responses to perturbations, even though its nonlinearity mirrored that of the minimal model where all reactions were cofactor-coupled.

The present study showed that there are similar metabolic responses to perturbations across three distinct kinetic models of *E*.*coli* metabolism. We like to underscore that these models were proposed by different research groups and within distinct contexts. Furthermore, the specifics of each model diverge considerably: both the Khodayari and Chassagnole models are tailored for aerobic conditions, whereas the Boecker model is for anaerobic conditions. Certain reactions and metabolites present in the Khodayari model are absent in both the Boecker and Chassagnole models. The reaction rate functions and parameter values differ across these models because each model has been developed to recapitulate different experimental results. Yet, in spite of these pronounced disparities, the models exhibit converging characteristics to a certain degree, notably the pivotal role of AXPs and the diminishing responsiveness attributed to the addition of reactions. Such congruence in outcomes might hint at the presence of features that remain resilient against variations in model specifics, potentially being shaped by the foundational network structures inherent to core metabolic processes.

From a theoretical standpoint, the robustness of metabolic systems, as well as the interplay between network structure and reaction dynamics, have been explored in the chemical reaction network theory and related fields Feinberg (2019); Gutenkunst et al. (2007); Himeoka et al. (2022b); Kobayashi et al. (2022). However, the practical relevance of these theories to metabolic systems remains to be ascertained since the theories are mathematically rigorous and the strong assumptions narrow down the applicable systems. The cross-talk between bacterial physiology and chemical reaction network theory should enhance the developments of each field.

In the following paragraphs, we discuss potential implications of strong responsiveness in the context of biological sciences. The unique feature of strong responsiveness, as observed in our realistic metabolic model, naturally prompts the question: what could be its biological function? An initial hypothesis suggests its role in sensing ambient condition changes and fluctuations. Wellknown design principles of signal transduction systems emphasize the amplification of external stimuli Ferrell and Ha (2014). Given that certain transcription factors are known to sense concentrations of metabolites in central carbon metabolism Kotte et al. (2010); Okano et al. (2019); Kochanowski et al. (2013), strong responsiveness could serve as an amplifier for external cues.

Exploring metabolic dynamics also holds promise for synthetic biology and metabolic engineering Shimizu et al. (2001); Schwander et al. (2016); Bar-Even et al. (2010); Doerr et al. (2019); Claassens et al. (2019); Kurisu et al. (2023); Opgenorth et al. (2017, 2014). The construction of artificial metabolic reaction networks often relies on pathway prediction algorithms, constraint-based models, and machine learning Moriya et al. (2010); Araki et al. (2015); Orth et al. (2010); Varma and Palsson (1993); Segler et al. (2018). While these algorithms consider stoichiometric feasibility, they frequently overlook dynamics. This oversight can result in proposed network structures underperforming Schwander et al. (2016); Bar-Even et al. (2010); Doerr et al. (2019); Claassens et al. (2019), especially in terms of cofactor recycling Opgenorth et al. (2017, 2014). Notably, *in silico* reconstructed metabolic systems often disrupt cofactor balance Okamoto et al. (1980); Okamoto and Hayashi (1983), posing challenges to bottom-up synthesis in artificial cells. We hope our study offers foundational insights for the development of quantitative metabolic dynamics theories.

## Acknowledgments

The authors thank Namiko Mitarai and Taro Furubayashi for fruitful discussions. This work is supported by JSPS KAKENHI Grant Numbers JP 21K20626, JP 22K15069, and JP 22H05403 to YH; JP 22K21344 to CF;, the Japan Science and Technology Agency (JST) ERATO (JPMJER1902 to CF), and GteX Program (JPMJGX23B4 to YH).

## Code availability

Codes for generating the computational models studied in the manuscript are deposited on GitHub (*https://github.com/yhimeoka/Perturbation-Response-Analysis.git*).

## Materials and methods

### Models and modifications

In Khodayari et al. Khodayari et al. (2014), the kinetic parameter values are estimated by using the ensemble modeling Khodayari et al. (2014); Tran et al. (2008); Tan and Liao (2012); Rizk and Liao (2009) so that the steady flux distribution of the metabolic reactions becomes consistent with the experimental measurements. However, the steady-state dealt with in the paper turned out to be unstable, i.e., the maximum eigenvalue of the Jacobi matrix of the linearized system at the steady state was positive. For carrying out the perturbation-response simulation, we need to make the steady-state stable. To make the steady-state stable, we fixed the periplasmic hydrogen concentration to the steady-state value and removed the competitive inhibition of phosphoenolpyruvate carboxylase (Those two factors were manually found). Also, we set the concentrations of the metabolites in the culture media (extracellular metabolites) with the assumption that the model cell is growing in a sufficiently large flask. In addition, the small molecules, H_2_O, O_2_, CO_2_, NH_4_ and phosphate, are set to be constant with further assuming that the exchange rate of the small chemicals between the extracellular culture is sufficiently fast.

For the dynamics of ATP and ADP concentrations, in the Chassagnole model Chassagnole et al. (2002), we used the reactions that coupled with ATP and ADP. Since the kinetic rate equations of those reactions are defined with the dependency of ATP and ADP concentrations in the original model, we utilized the kinetic rate equations as they are. The reactions for the adenine base synthesis are not modeled in the model, and thus, the total concentration of ATP and ADP are the conserved quantity in the model. By assuming that the decreases of the ATP and ADP due to the growth dilution are compensated, the growth dilution term is not introduced to the model equa-tions of ATP and ADP concentrations. Lastly, we introduced the non-growth associated ATP hydrolysis reaction (ATP→ADP) since the model fails to balance ATP and ADP and relaxes to a steady state where the concentration of some metabolites is almost zero. We utilized a simple MichaelisMenten rate equation for the hydrolysis reaction *J*_AH_ = *vx*_ATP_/(*K*_*M*_ + *x*_ATP_), where *v* = 0.1 (sec^−1^) and *K*_*M*_ = 1.0 (mM)

The Boecker model Boecker et al. (2021) is used without modification.

### Computation of dynamics

All the ODE simulations in the present manuscript are done using ode15s() function in MATLAB (MathWorks) Inc. (2022). For computing the attractor of the Chassagnole model, we used the initial concentrations registered in chassagnole4 on JWS online Olivier and Snoep (2004) as the initial concentrations of the modified Chassagnole model to run the dynamics. The steady-state concentrations of the metabolites reach from the initial concentrations and are utilized as the attractor.

Estimate of the fluctuation strength

For estimating the strength of fluctuations, we utilize the following simple metabolic reaction

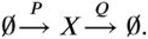

The concentration of the enzyme *P* and *Q* follows the stochastic transcription and translation model Friedman et al. (2006); Paulsson and Ehrenberg (2000). The temporal change of the concentration of the metabolite *X* here is given by a simple model with the mass-action kinetics,

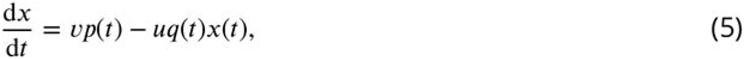

where *x, p, q* are the concentration of *X, P*, and *Q*, respectively. *v* and *u* are the kinetic parameters of the reactions. According to the time-scale separation of metabolic reactions and transcription/translation, we suppose that *x* is the fast variable and its dynamics slave to *p* and *q*. We assume that the timescales in the change of the metabolite concentrations are sufficiently smaller than that of the proteins and apply the quasi-steady-state approximation: we solve the steady-state of the deterministic equation of (5) Kaneko (1981); Risken (1996). Then, we calculate the mean and variance of the solution *x*(*t*) = *vp*(*t*)/*uq*(*t*), which is a stochastic variable. Since the stochastic transcription and translation model leads to the Gamma distribution of the protein concentration, the average and the variance are given by

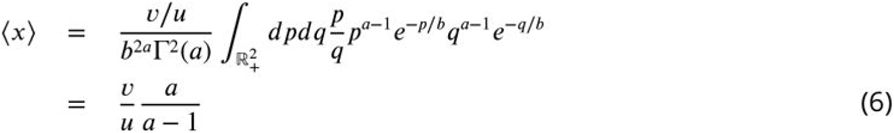

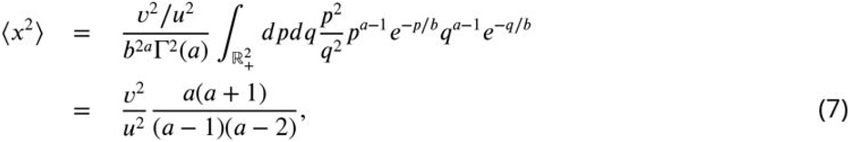

where *a* and *b* are the parameters set by the rate of the transcription, translation, mRNA degradation, and protein degradation.

The coefficient of variation is then given by

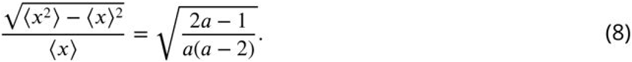

*a* is the ratio between the transcription rate and the protein degradation rate. According to the large-scale proteomic analysis of *E*.*coli* Taniguchi et al. (2010), the mean value of *a* among the essential genes is ≈ 6.82 ± 2.34 (only with the essential metabolic enzymes listed in the iML1515 Monk et al. (2017), the average value is ≈ 6.72 ± 2.34). By substituting *a* = 6.82 into (8), we obtain the relative noise strength as ≈ 62%. Since in the estimate, we suppose the irreversible reactions lead to larger noise amplitudes than reversible reactions, we regard 62% as the maximum relative noise strength and use a bit smaller value, 40%.

### The random catalytic reaction network model

The random catalytic reaction network (RCRN) model is a simple, mass-balancing toy model of cellular metabolism Furusawa and Kaneko (2003); Kaneko et al. (2015). The ordinary differential equation dictating the temporal evolution of the *i*th metabolite’s concentration *x*_*i*_ is given by

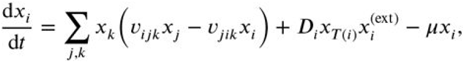

where *v*_*ijk*_, *D*_*i*_ and *μ* are the rate constant of the reaction C_*j*_ → C_*i*_ catalyzed by C_*k*_ (C_*i*_ denotes the *i*th metabolite), the substrate update rate of C_*i*_, and the specific growth rate of the model cell, respectively. *T* (*i*) denotes the transporter metabolite (enzyme) for the uptake of C_*i*_, and 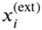 is the external concentration of C_*i*_. In the RCRN model, the total volume of the cell is usually set to be equal to the total amount of the metabolites. This is because each “metabolite” in the model is interpreted as an enzyme as well as a reactant of the metabolic reactions. Hence, the specific growth rate of the model cell is set to the total uptake rate of the metabolites, 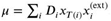.

In the construction of an instance of the RCRN model, we first generate a random, connected network among *N* metabolites, and then, we assign a single catalyst on each reaction edge. If the reaction C_*j*_ → C_*i*_ catalyzed by C_*k*_ exists in the network, *v*_*ijk*_ is non-zero, while otherwise, it is set to zero. In the present manuscript, we suppose all the reactions are reversible, and thus, *v*_*jik*_ is non-zero if and only if *v*_*ijk*_ is non-zero.

For making the comparison of the responsiveness of the realistic model to the RCRN model, we generated 128 instances of the RCRN model where the number of metabolites *N* is set to be equal to the realistic model that we want to make a comparison. The number of reactions *R* is set to 3.5*N* for the Boecker and Chassagnole model while 4*N* for the Khodayari model. The ratio between *N* and *R* is set so that the Erdős–Rényi random graph of given size becomes connected with sufficiently high probability. The reaction rate constants *v*_*ijk*_’s are sampled from the model to compare. For the sake of simplicity, we suppose only the 1st metabolite is the nutrient metabolite which is taken up from the external environment and the *N*th metabolite is the transporter of it.

*D*_*i*_ and 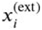 are set to unity for *i* = 1, and for the else, those parameters are set to zero.

For each generated instance of the RCRN model, we performed the perturbation response simulation where the attractor(s) are computed by simulating the model dynamics from randomly generated 32 initial concentrations ***x***_ini_ ∈ (10^−3^, 10^3^)^*N*^. The RCRN models exhibiting multistability or computational failure due to numerical issues such as heavy stiffness are not subjected to further analysis (Less than 10% of the model instances showed multistability for corresponding models. The computational failure is ≈ 20% for the RCRN model with the parameter distribution of Chassagnole model, ≈ 50% for that of the Khodayari model, while less than 1% for that of the Boecker model). For quantifying the response coefficient, we computed 128 trajectories for each of those models.

### Random addition of the metabolic reactions

First, we have add uni-uni reactions where the reactants are randomly picked from the non-cofactor metabolites. Next, we decide whether the additional reaction to be coupled with the adenyl cofactor metabolites (ATP, ADP, or AMP) with a propability *p*. The probability *p* equals to the fraction of adenyl cofactor-coupled reactions in the original model. The addition reaction is either the uni-uni reaction (if the reaction is not coupled with AXPs) or the bi-bi reaction (if the reaction is coupled with the adenyl cofactors). We computed the parameter value distribution for each reaction scheme and the parameter values for the additional reactions are sampled from the corresponding parameter value distribution.

For the Khodayari model, the additional reaction is modeled using the elementary reaction decomposition (see the next section), i.e, a reaction *A* ⇌ *B* is decomposed into the three elementary reaction steps, *E* + *A* ⇌ *EA, EA* ⇌ *EB*, and *EB* ⇌ *E* + *B*. The mass-action kinetics is adopted as the reaction rate function for each elementary reaction step.

The elementary reaction decomposition is not applied to the Boecker model and the Chassagnole model. Also, several types of reaction rate functions are used in the two models. Thus, we decided to use the simplest enzymatic reaction kinetics used in each paper. The reversible Michaelis-Menten kinetics

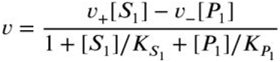

is used for uni-uni reactions, and for bi-bi reactions, the ordered bi-bi reaction kinetics

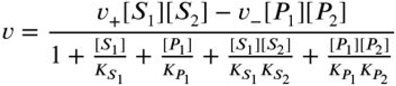

is adopted in the Boecker model, a reaction kinetics

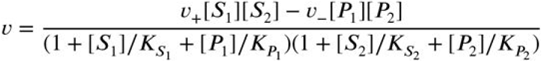

is used in the Chassagnole model. [*S*_*i*_], [*P*_*i*_], and *K*_*_ (* is either *S*_1_, *S*_2_, *P*_1_ or *P*_2_) represent the concentration of the *i*th substrate and product, and the dissociation constant of the corresponding chemical, respectively.

After constructing an extended model, we compute the steady-state attractor by simulating the dynamics from 128 random initial concentrations. These initial conditions are generated by applying a 40% relative perturbation to the steady-state concentrations of the original model. If all the initial conditions converge to a single attractor, then we generate another set of 128 initial conditions by applying a 40% relative perturbation to the steady-state concentrations of the extended model in order to perform the perturbation-response simulation. For the Boecker model, we require the extended model to have a steady growth rate greater than half of that of the original model. This is because growth dilution is typically the slowest process in the model, and thus, a significant slowdown of growth dilution can lead to an artificial overestimation of the model’s responsiveness.

### Elementary reaction decomposition

Here, a brief description of the elementary reaction decomposition (ERD) is provided. Suppose that we have the following bi-to-uni reaction

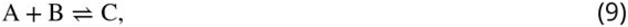

and that the reaction is catalyzed by an enzyme E. For implementing this reaction into the kinetic model, the Michaelis-Menten type kinetics of chemical reaction is often adopted. With the Michaelis-Menten type kinetics, the rate of the reaction *J* is given by

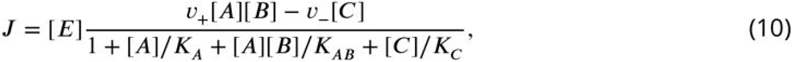

where we supposed that the binding of the substrate to the enzyme occurs in the order of *A* → *B*.

The Michaelis-Meneten type reaction kinetics ((10)) is obtained by the pseudo-equilibrium approximation or the steady-state approximation of the following reactions

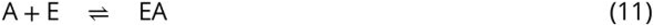

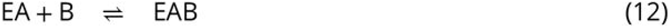

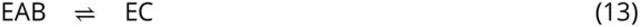

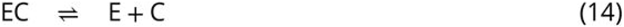

In the ERD scheme, each step ((11)-(14)) of the overall reaction ((9)) is modeled using the massaction kinetics. Also, the forward- and the backward reactions are dealt as different reactions. For instance, the rates of the reaction A + E ⇌ EA is given by *J*_*A,E*→*EA*_ = *v*_*A,E*→*EA*_[*E*][*A*] and *J*_*EA*→*A,E*_ = *v*_*EA*→*A,E*_[*EA*].

The advantage of the ERD is that every reaction rate function is given either by a linear or quadratic function of the metabolites’ concentrations. This feature allows us, in the present manuscript, to randomize the reaction network in a unified manner. Also, the ERD has the advantage of reduc-ing the computational cost for the parameter estimation based on omics data. For more details, see original papers Khodayari et al. (2014); Tran et al. (2008); Tan and Liao (2012); Rizk and Liao (2009).

### The generative AIs and other software

ChatGPT (OpenAI, September 25, 2023 version) and Grammarly (Grammarly, Inc.) are used to improve clarity and to brush up on English grammar.

## Supplementary tables and figures

**Table S1.** The list of the substrate-level regulations in the Khodayari model.

| Reaction ID | Metabolites ID | Type of regulation |
| --- | --- | --- |
| ACALD | amp | inhibition |
| FBA | dhap,adp,amp,pep,pi | inhibition |
| FUM | cit | inhibition |
| G6PDH2r | nadh,nadph | inhibition |
| GLUN | adp,atp,glu_L,nad,nadh,nadp,nadph,pi | inhibition |
| GLUSy | glu_L | inhibition |
| GND | atp,fdp,nadp,ru5p_D | inhibition |
| ICDHyr | glx,oaa,pep | inhibition |
| ICL | akg,icit,prp,succ | inhibition |
| LDH | atp | inhibition |
| PGI | 6pgc,pep | inhibition |
| PPC | mal_L | inhibition |
| PPCK | atp,pep | inhibition |
| PPS | adp,akg,amp,oaa,pep | inhibition |
| PTAr | adp,atp,nadh,nadph,pep,pyr | inhibition |
| PYKF | atp,succoa | inhibition |
| RPI | amp | inhibition |
| TALA | pi | inhibition |
| PYKA | atp,succoa | inhibition |
| EDA | 6pgc,g3p | inhibition |
| PGMT | accoa | inhibition |
| GLGC | amp | inhibition |

**Table S2.** The list of the substrate-level regulations in the Boecker model.

| Reaction ID | Metabolites ID | Type of regulation |
| --- | --- | --- |
| PFK | ADP,FBP | activation |
| PTACK | PYR,PEP | activation |
| FK | PEP | inhibition |
| PYK | ATP,PYR | inhibition |
| CSICD | AKG,NADH | inhibition |
| PTACK | ATP | inhibition |
| PPC | FUM | inhibition |
| PFL | AcCoA,PYR | inhibition |
| ADH | ACTLD,NAD | inhibition |
| MDH | OAA,NAD,NADH,FUM | inhibition |
| FRD | OAA | inhibition |
| ALDH | CoA | inhibition |

**Table S3.** The list of the substrate-level regulations in the Chassagnole model.

| Reaction ID | Metabolites ID | Type of regulation |
| --- | --- | --- |
| PTS | G6P | inhibition |
| PGDH | atp | inhibition |
| PGL | 6pg | inhibition |
| G1PAT | fdp | activation |
| PFK | pep | inhibition |
| PFK | adp | activation |
| PEPCxylase | fdp | activation |
| PK | fdp | activation |
| PK | atp | inhibition |

**Table S4.**
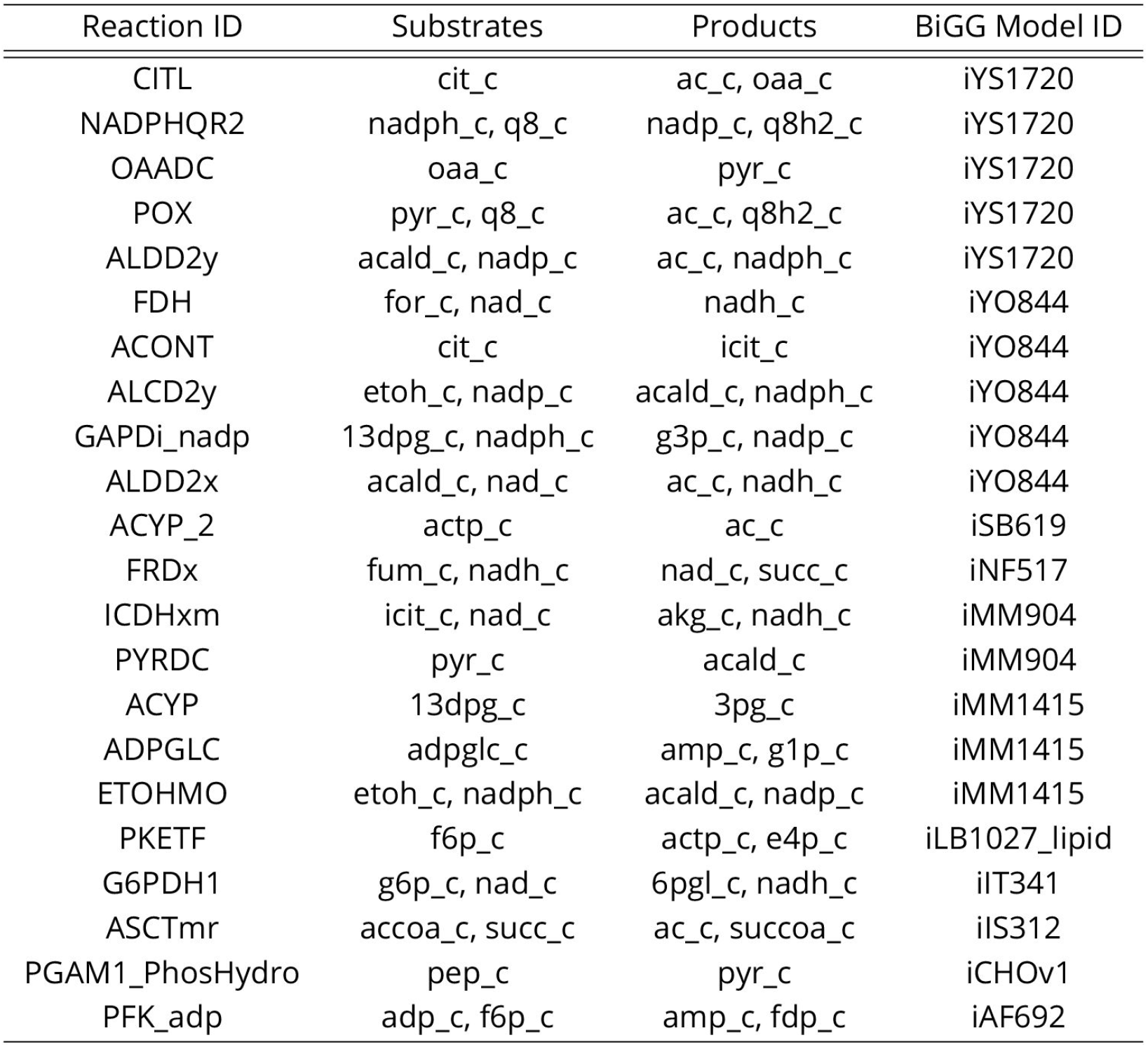
The list of the additional reactions used for generating Fig. S7.

**Figure S1.**
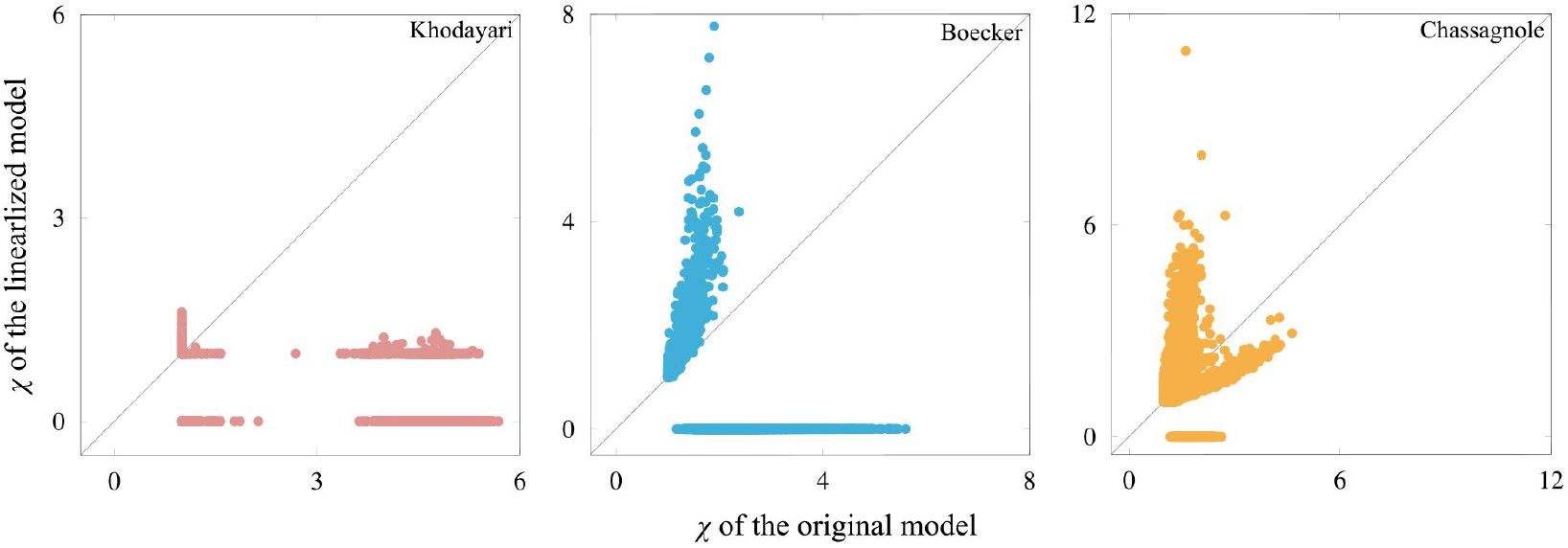
Scatter plot of the response coefficient. Each point corresponds to a single initial condition, while the horizontal and vertical axes represent the response coefficient *χ* computed by the original and the linearized model, respectively, around the steady state. The linearized model is given by *d****x***/*dt* = *J* ***x***, where *J* is the Jacobi matrix of the model at steady state. If any metabolite concentrations become negative, the response coefficient is incomputable and “0” is assigned to such trajectories (note that the minimum value of the response coefficient is unity). Since the response coefficient is based on the logarithm of the concentrations, as the metabolite concentrations approach zero, the response coefficient becomes larger. The high response coefficient in the Boecker and Chassagnole model would be explained by this artifact. The linearized Khodayari model shows either *χ* ≈ 1 or *χ* = 0 (one or more metabolite concentrations become negative).This could be due to the number of variables in the model. For the response coefficient to have a larger value, the perturbation should be along the eigenvector that leads to oscillatory dynamics with long relaxation time (i.e., the corresponding eigenvalue has a small real part in terms of absolute value and a non-zero imaginary part). However, since the Khodayari model has about 800 variables, if perturbations are along such directions, there is a high probability that one or more metabolite concentrations will become negative.

**Figure S2.**
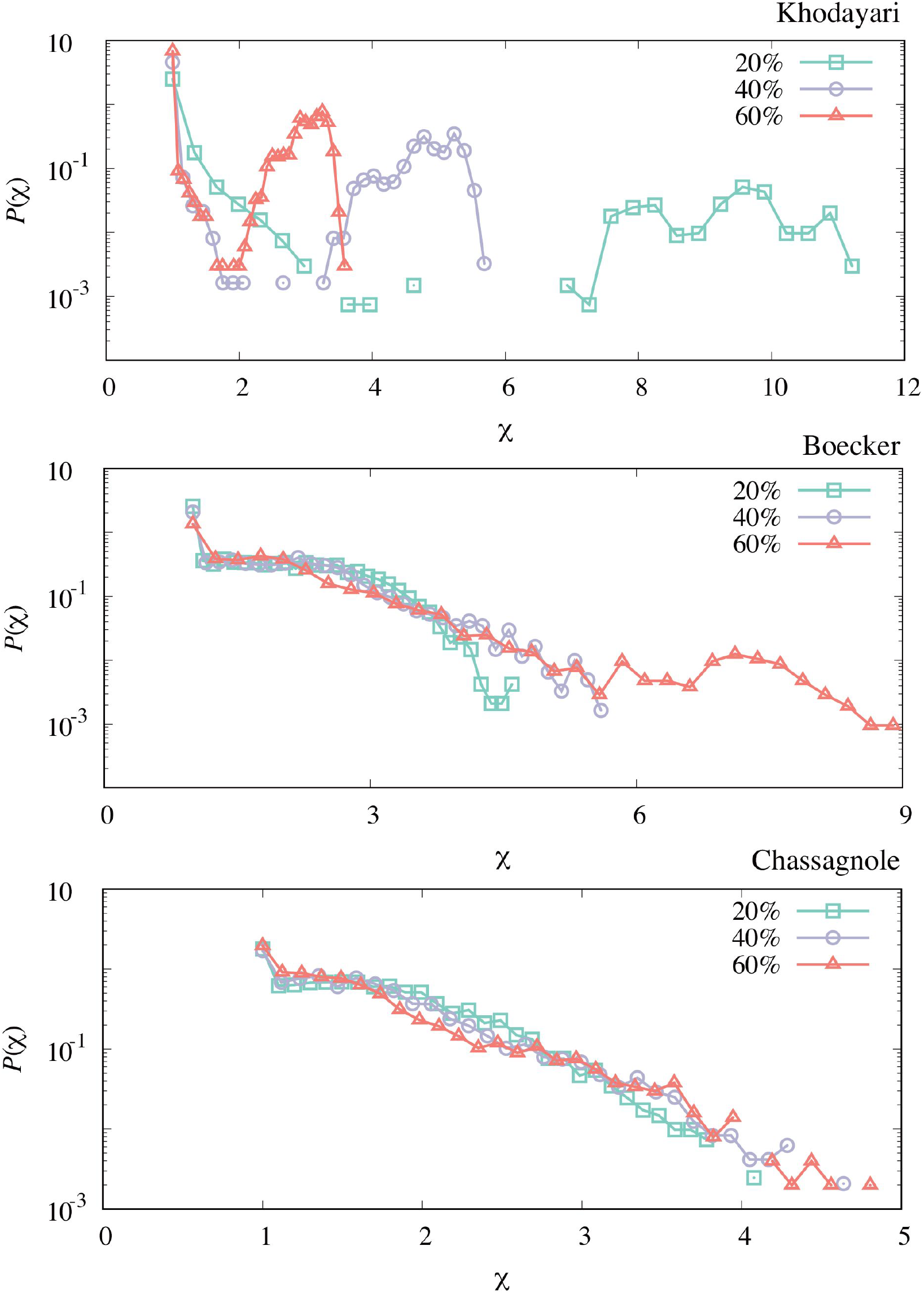
The response coefficient distribution of each model obtained by the perturbation-response simulation with different perturbation strengths (20%, 40% and 60% relative perturbation). Each distribution is obtained by simulating 4096 trajectories. The qualitative features (double-peaks for the Khodayari model and the plateau for the Boecker and Chassagnole models) are unvaried, while the scales of the response coefficient of the Khodayari model differ among the perturbation strengths.

**Figure S3.**
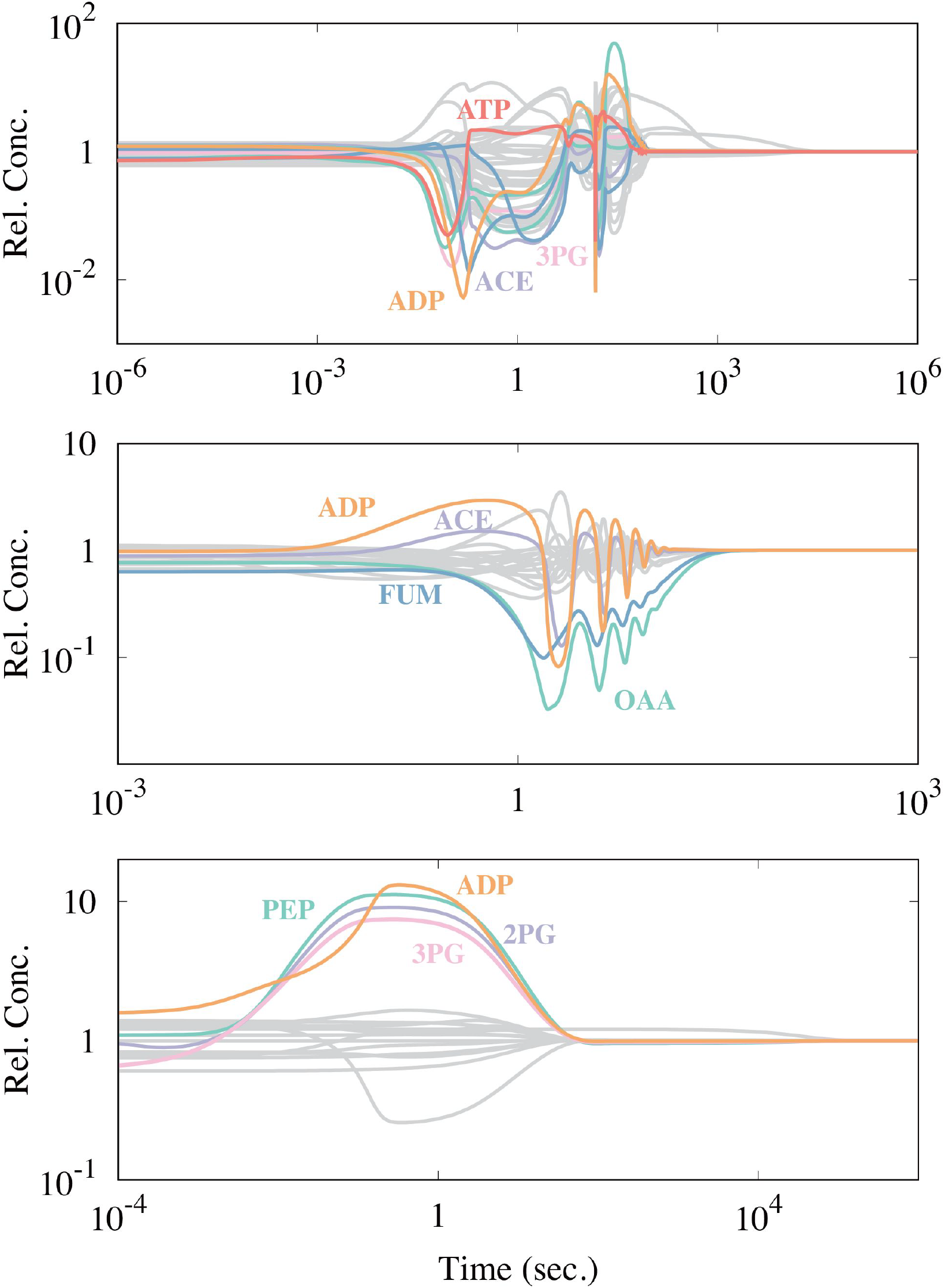
The time course of each model with the strongly responding metabolites being highlighted. A metabolite is highlighted if its concentration changes more than *X*-fold from its steady concentration. *X* is set to be 5 for the Boecker and Chassagnole models, while *X* = 25 is chosen for the Khodayari model because too many metabolites are highlighted with *X* = 5.

**Figure S4.**
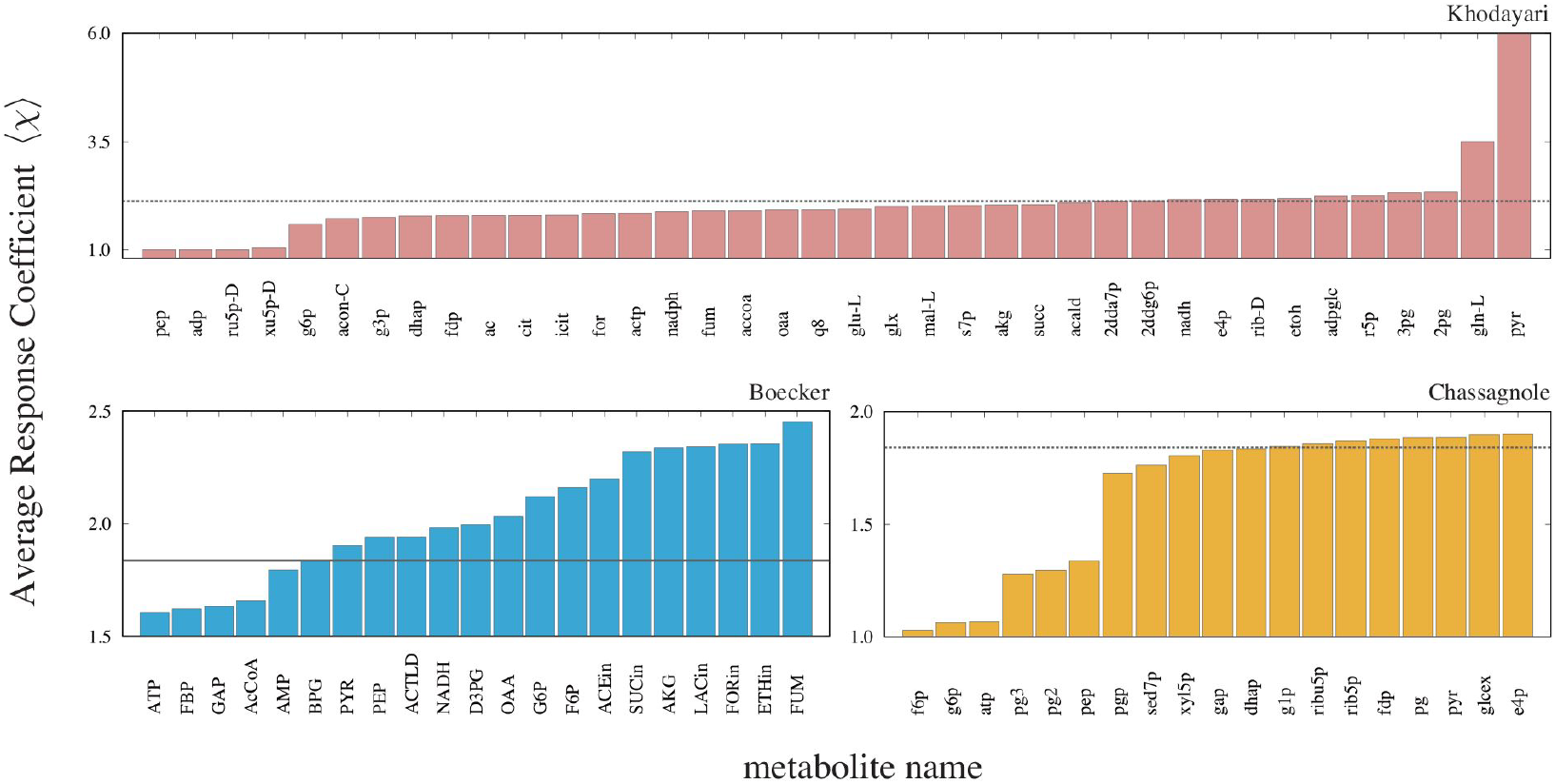
The average response coefficients of the models with the concentration of indicated metabolite fixed. The dashed line in each panel is the average response coefficient of the original model.

**Figure S5.**
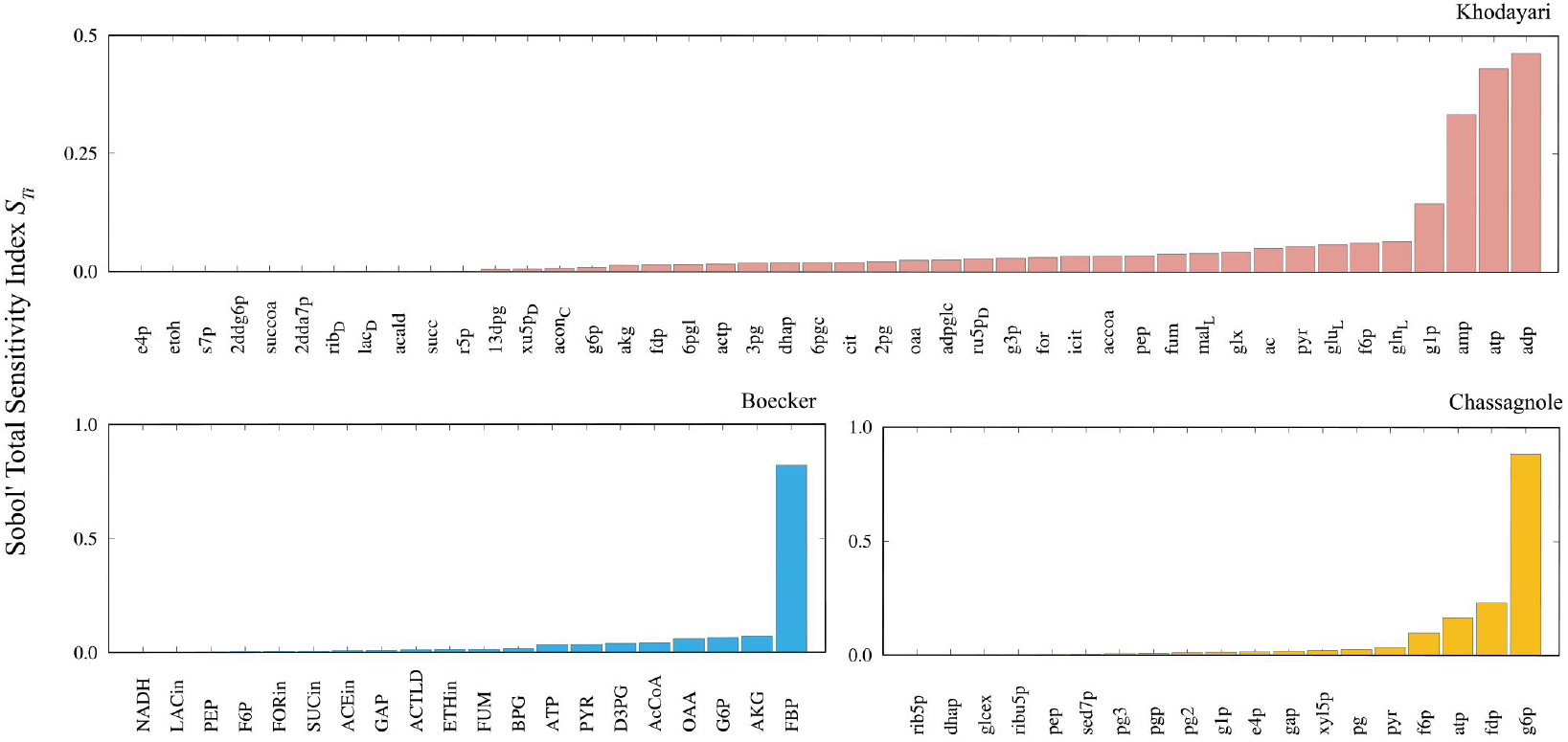
The Sobol’ total sensitivity index 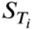 is computed for each metabolite. The sensitivity indices are computed by simulating 512 trajectories for the Khodayari model and 4096 trajectories for the Boecker and Chassagnole models.

**Figure S6.**
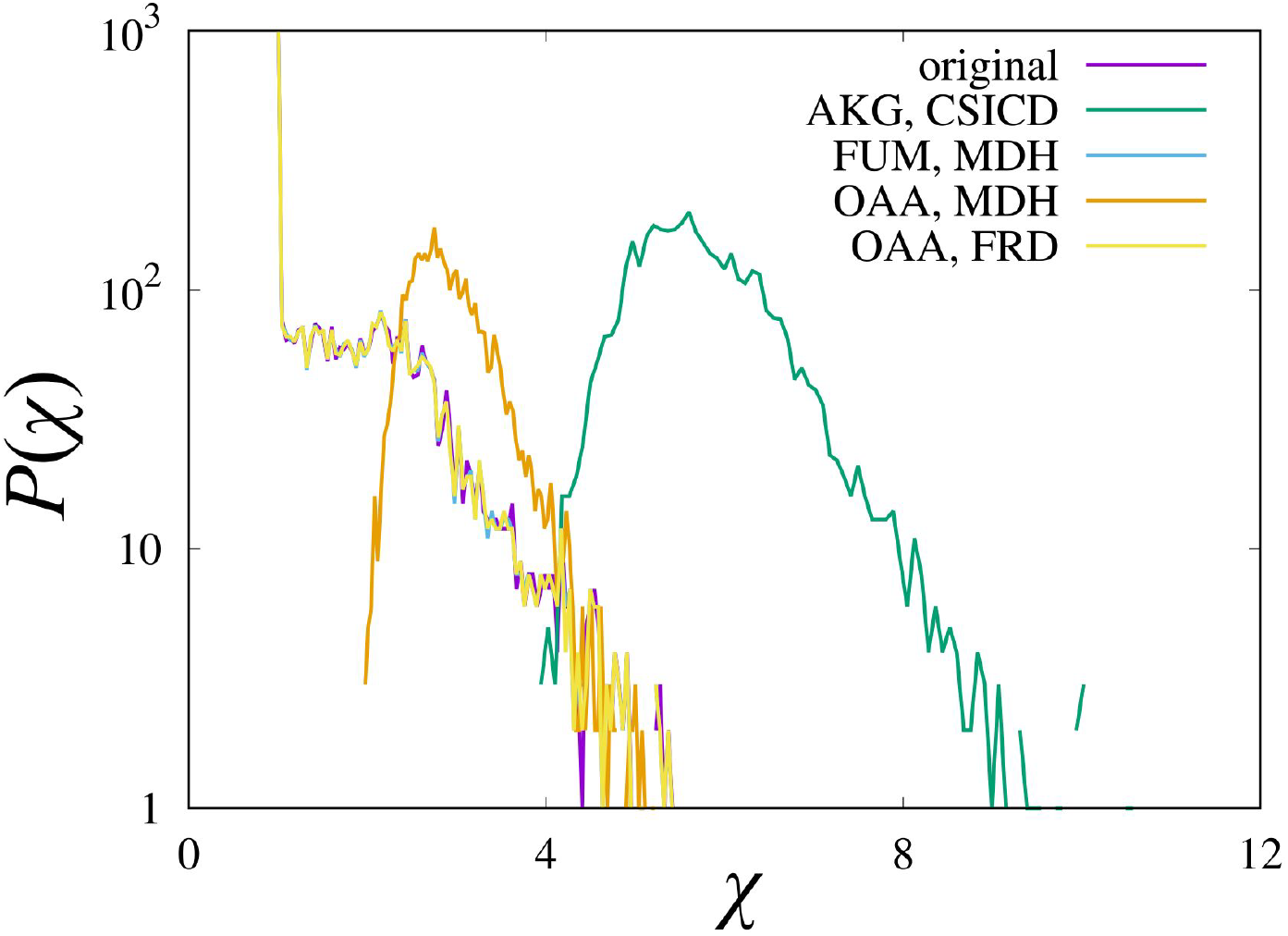
The response coefficient distributions of models with weakened substrate-level regulation of enzyme activity. The relaxations of the inhibition strength of AKG on CSICD and OAA on MDH increase the responsiveness of the model, while those of FUM on MDH and OAA on FRD have no effect. To relax the inhibition, the equilibrium constant for the corresponding competitive inhibition *k*_rxn_i_met_ is set to 10^6^ × *k*_rxn_i_met_ (the inhibition term is implemented with the form *x*_met_ /*k*_rxn_i_met_ in the denominator of the reaction rate function), where rxn, i, and met represent the reaction name, “inhibition”, and the metabolite name, respectively. Each distribution is computed from 4096 trajectories.

**Figure S7.**
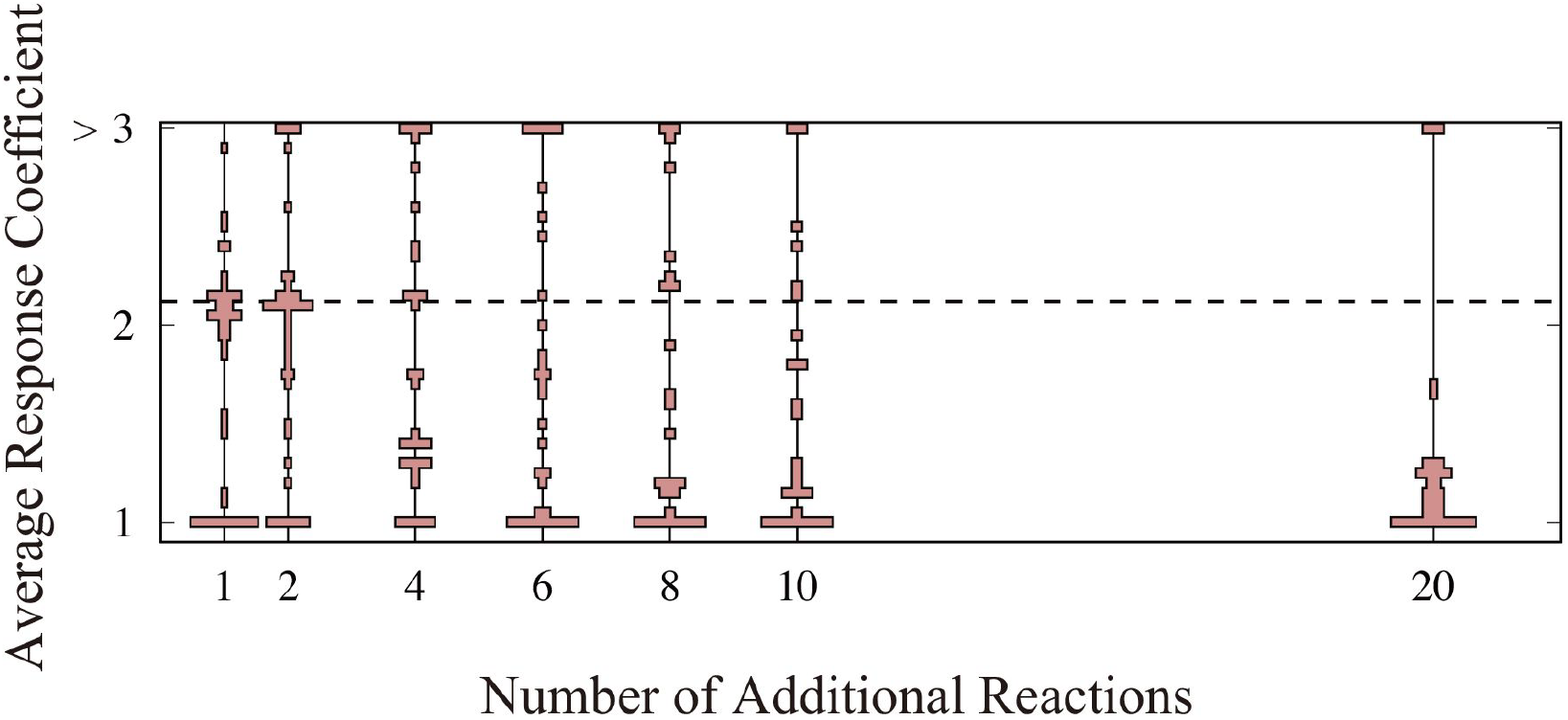
The violin plot of the average response coefficient of the network expansion of the Khodayari model. The horizontal axis represents the number of additional reactions. The additional reactions are chosen from the reactions registered in the metabolic model database (see **Table. S4**).

**Figure S8.**
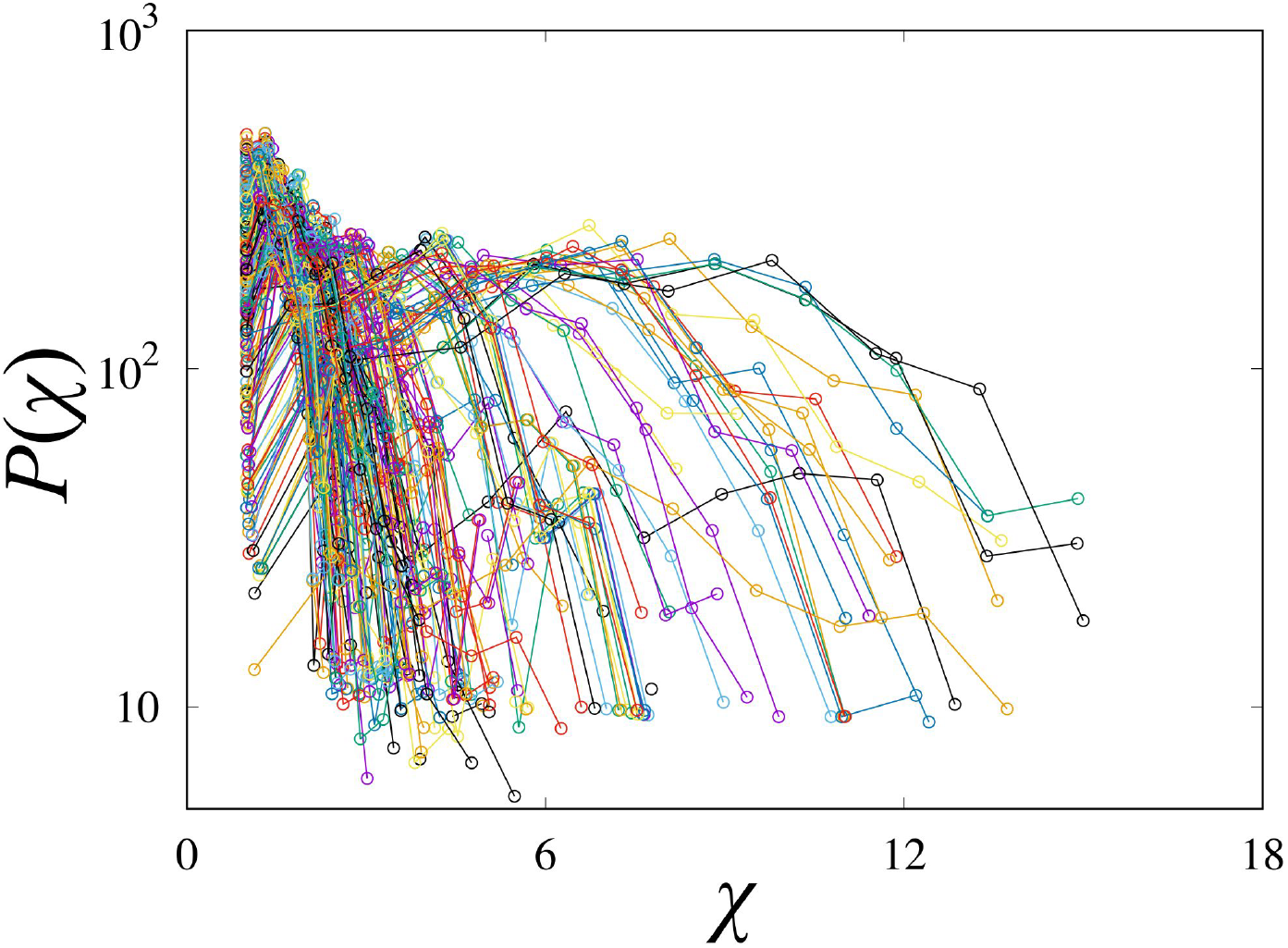
The response coefficient distribution of 215 realizations of the PYK model by the MASSpy framework Haiman et al. (2021) are overlaied (each model has the same network structure, kinetic rate law, but the parameters are different). For each model realization, we have computed 256 trajectories. Trajectories had negative value for any metabolites are excluded for the computation of the distribution.

## Editors

### Reviewing Editor

**Marisa Nicolás**

Laboratório Nacional de Computação Científica, Rio de Janeiro, Brazil

### Senior Editor

**Aleksandra Walczak**

École Normale Supérieure - PSL, Paris, France

#### Reviewer #1 (Public review)

(1a) Summary:

The author studied metabolic networks for central metabolism, focusing on how system trajectories returned to their steady state. To quantify the response, systematic perturbation was performed in simulation and the maximal destabilization away from steady state (compared with initial perturbation distance) was characterized. The author analyzed the perturbation response and found that sparse network and networks with more cofactors are more “stable”, in the sense that the perturbed trajectories have smaller deviation along the path back to the steady state.

(1b) Strengths and major contributions:

The author compared three metabolic models and performed systematic perturbation analysis in simulation. This is the first work characterized how perturbed trajectories deviate from equilibrium in large biochemical systems and illustrated interesting findings about the difference between sparse biological systems and randomly simulated reaction networks.

(1c) Weaknesses:

There are two main weaknesses in this study:

First, the metabolic network in this study is incomplete. For example, amino acid synthesis and lipid synthesis are important for biomass and growth, but they are not included in the three models used in this study. NADH and NADPH are as important as ATP/ADP/AMP, but they are not included in the models. In the future, a more comprehensive metabolic and biosynthesis model is required.

Second, this work does not provide mathematics explanation on the perturbation response χ. Since the perturbation analysis are performed closed to steady state (or at least belongs to the attractor of single steady state), local linear analysis would provide useful information. By complement with other analysis in dynamical systems (described in below) we can gain more logical insights about perturbation response.

(1d) Discussion and impact for the field:

Metabolic perturbation is an important topic in cell biology and has important clinical implication in pharmacodynamics. The computational analysis in this study provides an initiative for future quantitative analysis on metabolism and homeostasis.

Comments on revised version:

The revised version of this manuscript made some clarifications, while I think the analysis of response coefficients is still numerical and model-specific, being unclear under dynamical systems of views.

https://doi.org/10.7554/eLife.98800.2.sa2

#### Reviewer #2 (Public review)

The authors have conducted a valuable comparative analysis of perturbation responses in three nonlinear kinetic models of E. coli central carbon metabolism found in the literature. They aimed to uncover commonalities and emergent properties in the perturbation responses of bacterial metabolism. They discovered that perturbations in the initial concentrations of specific metabolites, such as adenylate cofactors and pyruvate, significantly affect the maximal deviation of the responses from steady-state values. Furthermore, they explored whether the network connectivity (sparse versus dense connections) influences these perturbation responses. The manuscript is reasonably well written.

Comments on revised version:

The authors have addressed my concerns to a large extent. However, a few minor issues remain, as listed below:

1. The authors identified key metabolites affecting responses to perturbations in two ways: by fixing a metabolite’s value and (ii) by performing a sensitivity analysis. It would be helpful for the modeling community to understand better the differences and similarities in the obtained results. Do both methods identify substrate-level regulators? Is freezing a metabolite’s dynamics dramatically changing the metabolic response (and if yes, which ones are so different in the two cases)? Does the scope of the network affect these differences and similarities?
2. Regarding the issues the authors encountered when performing the sensitivity analysis, they can be approached in two ways. First, the authors can check the methods for computing conserved moieties nicely explained by Sauro’s group (doi:10.1093/bioinformatics/bti800) and compute them for large-scale networks (but beware of metabolites that belong to several conserved pools). Otherwise, the conserved pools of metabolites can be considered as variables in the sensitivity analysis-grouping multiple parameters is a common approach in sensitivity analysis. https://doi.org/10.7554/eLife.98800.2.sa1

### Author response

The following is the authors’ response to the original reviews.

***Reviewer #1 (Public Reviews):***

*First, the metabolic network in this study is incomplete. For example, amino acid synthesis and lipid synthesis are important for biomass and growth, but they4 are not included in the three models used in this study. NADH and NADPH are as important as ATP/ADP/AMP, but they are not included in the models. In the future, a more comprehensive metabolic and biosynthesis model is required*.

Thank you for the critical comment on the weakness of the present study. We actually tried to study a larger model like Turnborg et al (2021), which is a model of JCVI-syn3A, but we give up to include it in our model list to study in depth. This is because we noticed that the concentration of ATP in the model can be negative (we confirmed this with one of the authors of the paper). Another “big” kinetic model of metabolism that we could list would be Khodayari et al (2017). However, we could not find the models to compare the dynamics of this big model with. Therefore, we decided to use the model only for the central carbon metabolism for now. We would like to leave a more extended study for the near future.

We would like to mention that NADH and NADPH are included in Khodayari model and Boecker model, while NADH and NADPH are ramped up to NADH in the latter model.

> *Second, this work does not provide a mathematical explanation of the perturbation response χ. Since the perturbation analysis is performed close to the steady state (or at least belongs to the attractor of single-steady-state), local linear analysis would provide useful information. By complementing with other analysis in dynamical systems (described below) we can gain more logical insights about perturbation response*.

We tried a linear stability analysis. However, with the perturbation strength we used here, the linearization of the model is no longer valid, in the sense that the linearized model

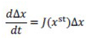

leads to negative concentrations of the metabolites (xst+Δx < 0 for some metabolites). We have added a scatter plot of the response coefficient of trajectories sharing the initial condition, while the dynamics are computed by the original model and the linearized model, respectively. (Fig. S1).

Since the response coefficient is based on the logarithm of the concentrations, as the metabolite concentrations approach zero, the response coefficient becomes larger. The high response coefficient in the Boecker and Chassagnole model would be explained by this artifact. The linearized Khodayari model shows either χ~1 or χ = 0 (one or more metabolite concentrations become negative). This could be due to the number of variables in the model. For the response coefficient to have a larger value, the perturbation should be along the eigenvector that leads to oscillatory dynamics with long relaxation time (i.e., the corresponding eigenvalue has a small real part in terms of absolute value and a non-zero imaginary part). However, since the Khodayari model has about 800 variables, if perturbations are along such directions, there is a high probability that one or more metabolite concentrations will become negative.

We fully agree that if the perturbations on the metabolite concentrations are in the linear regime, the response to the perturbations can be estimated by checking the eigenvalues and eigenvectors. However, we would say that the relationship between the linearized model (and thus the spectrum of eigenvalues) and the original model is unclear in this regime. We remarked this in Lines 158160.

> ***Recommendations for the authors:***
>
> *My major suggestion is about understanding the key quantity in this study: the response coefficient χ. When the perturbed state is close to the fixed point, one could adopt local stability analysis and consider the linearized system. For a linear system with one stable fixed point P, we consider the Jacobian matrix M on P. If all eigenvalues of M are real and negative, the perturbed trajectory will return to P with each component monotonically varies. If some eigenvalues have negative real part and nonzero imaginary part, then the perturbed trajectory will spiral inward to the fixed point. Depending on the spiral trajectory and the initially perturbed state, some components would deviate furthermore (transiently) from the fixed point on the spiral trajectory. This explains why the response coefficient χ can be greater than 1*.
>
> *Mathematically, a locally linearized system has similar behavior to the linear system, and the examples in this study can be analyzed in the similar way. Specifically, if a system has many complex eigenvalues, then the perturbed trajectory is more likely to have further deviation. The metabolic network models investigated in this work are not extremely large, and hence the author could analyze its spectrum of the Jacobian matrix at the steady state. Since the steady state is stable, I expect the spectrum located in the left half of the complex plane. If the spectrum spread out away from the real axis, we expect to see more spiral trajectories under perturbation. I think the spectrum analysis will provide a complementary view with respect to analysis on χ. The authors’ major findings, about the network sparsity and cofactors, can also be investigated under the framework of the spectrum analysis*.
>
> *Of course, when the nonlinear system is perturbed far away from the fixed point, there are other geometrical properties of the vector field that can cause the response coefficient χ to be greater than 1. This could also be investigated in the future by testing the behavior of small and large perturbations and observing if the systems have signatures of nonlinearity*.

Since all perturbed states return to the steady state, the eigenvalues of the Jacobi matrix accompanying the linearized system around the steady state are in the left half complex plane (negative real value). Also, some eigenvalues have non-zero imaginary parts.

The reason we emphasize the “nonlinear regime” is that the linearization is no longer valid in this regime, i.e. the metabolite concentrations can be negative when we calculate the linearized system. Certainly, there are complex eigenvalues in the Jacobi matrix of any model. However, we would say that there is no clear relationship between the eigenvalues and the response coefficient.

> *Minor suggestions:*
>
> *Line 127: Regarding the source of perturbation, cell division also generates unequal concentration of proteins and metabolites for two daughter cells, and it is an interesting mechanism to create metabolic perturbation*.
>
> Thank you for the insightful suggestion. We mentioned the cell division as another source of perturbation (Lines 130-131).
>
> *Line 175: I do not quite understand the statement “fixing each metabolite concentration*…*”, since the metabolite concentration in the ODE simulation would change immediately after this fixing*.

We meant in the sentence that we fixed the concentration of the selected metabolite as the steady state concentration and set the dx/dt of that metabolite to zero. We have rewritten the sentences to avoid confusion (Lines 180-181).

> *Figure 2: There are a lot of inconsistencies between the three models. Could we learn which model is more reasonable, or the conclusion here is that the cellular response under perturbation is model-specific? The latter explanation may not be quite satisfactory since we expect the overall cellular property should not be sensitive to the model details*.

Ideally, the overall cellular property should be insensitive to model details. However, the reality is that the behavior of the models (e.g., steady-state properties, relaxation dynamics, etc.) depends on the specific parameter choices, including what regulation is implemented. I think this situation is part of the motivation for the ensemble modeling (by J. Liao and colleague) that has been developed.

Detailed responsiveness would be model specific. For example, FBP has a fairly strong effect in the Boecker model, but less so in the Khodayari model, and the opposite effect in the Chassagnole model (Fig. 2). Our question was whether there are common tendencies among kinetic models that tend to show model-specific behavior.

> ***Reviewer 2 (Public Review):***
>
> *(1) In the study on determining key metabolites affecting responses to perturbations (starting from line 171), the authors fix the values of individual concentrations to their steady-state values and observe the responses. Such a procedure adds artificial constraints to the network because, in the natural responses of cells (and models) to perturbations, it is highly unlikely that metabolites will not evolve in time. By fixing the values of specific metabolites, the authors prohibit the metabolic network from evolving in the most optimal way to compensate for the perturbation. Instead of this procedure, have the authors considered for this task applying techniques from variance-based sensitivity analysis (Sobol, global sensitivity analysis), where they can calculate the first-order sensitivity index and total effect index? Using this technique, the authors would be able to determine the key metabolites while allowing for metabolic responses to perturbations without unnatural constraints*.

Thank you for the useful suggestion for studying the roles of each metabolite for responsiveness. We have computed the total sensitivity index (Homma and Salteli, 1996) for each metabolite of each model (Fig.S5). The total sensitivity indices of ATP are high-ranked in Khodayari- and Chassagnole model, while it is middle-ranked in the Boecker model. We believe that the importance of the adenyl cofactors is highlighted also in terms of the Sobol’ sensitivity analysis (the figure is referred in Lines 193-195).

We have encountered a minor difficulty for computing the sensitivity index. For the computation of the sensitivity index, we need to carry out the following Monte Carlo integral,

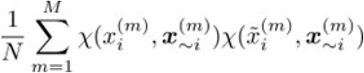

where the superscript (m) is the sample number index. The subscript i represents the ith element of the vector x, and ~i represents the vector x except for the ith element. The tilde stands for resampling.

There are several conserved quantities in each model. For independent resampling, we need to deal with the conserved quantities. For the Boecker and Chassagnole models, we picked a single metabolite from each conservation law and solved its concentration algebraically to make the metabolite concentration the dependent variable. Then, we can resample the metabolite concentration of one metabolite without changing the concentrations of other metabolites, which are independent variables.

However, in the Khodayari model, it was difficult to solve the dependent variables because the model has about 800 variables. Therefore, we gave up the computations of the sensitivity indices of the metabolites whose concentration is part of any conserved quantities, namely NAD, NADH, NADP, NADPH, Q8, and Q8H2.

> *(2) To follow up on the previous remark, the authors state that the metabolites that augment the response coefficient when their concentration is fixed tend to be allosteric regulators. The authors should report which allosteric regulations are implemented in each of the models so that one can compare against Figure 2. Again, the effect of allosteric regulation by a specific metabolite that is quantified the way the authors did is biased by fixing the concentration value - it is true that negative feedback is broken when the metabolite concentration is fixed, however, in the rate law, there is still the fixed inhibition term with its value corresponding to the inhibition at the steady state. To see the effect of allosteric regulation by a metabolite, one can change the inhibition constants instead of constraining the responses with fixed concentrations*.

We have listed the substrate-level regulations (Table S1-3). Also, we re-ran the simulation with reduced the effect of the substrate-level regulations for the reactions that are suspected to influence the change of the response coefficient. Instead of fixing the concentrations (Fig. S6).

The impact of substrate-level regulations is discussed in Lines 203-212.

We replaced “allosteric regulation” with “substrate-level regulation” because we noticed that some regulations are not necessarily allosteric.

> *(3) Given the role of ATP in metabolic processes, the authors’ finding of the sensitivity of the three networks’ responses to perturbations in the AXP concentrations seems reasonable. However, drawing such firm conclusions from only three models, with each of them built around one steady state and having one kinetic parameter set despite that they were built for different physiologies, raises some questions. It is well-known in studies related to basins of attraction of the steady states that the nonlinear responses also depend on the actual steady states, the values of kinetic parameters, and implemented kinetic rate law, i*.*e*., *not only on the topology of the underlying systems. In the population of only three models, we cannot exclude the possibility of overlaps and strong similarities in the values of kinetic parameters, steady states, and enzyme saturations that all affect and might bias the observed responses. Ideally, to eliminate the possibility of such biases, one should simulate responses of a large population of models for multiple physiologies (and the corresponding steady states) and multiple parameter sets per physiology. This can be a difficult task, but having more kinetic models in this work would go a long way toward more convincing results. Recently, E. coli nonlinear kinetic models from several groups appeared that might help in this task, e*.*g*., *Haiman et al*., *PLoS Comput Biol, 17(1): e1008208, (2021), Choudhury et al*., *Nat Mach Intell, 4, 710-719, (2022); Hu et al*., *Metab Eng, 82, 123-133 (2024), Narayanan et al*., *Nat Commun, 15:723, (2024)*.

We have computed the responsiveness of 215 models generated by the MASSpy package (Haiman et al, 2021). Several model realizations showed a strong responsiveness, i.e. a broader distribution of the response coefficient (Fig.S8), and mentioned in Lines 339-341.

We would like to mention that the three models studied in the present manuscript have limited overlap in terms of kinetic rate law and, accordingly, parameter values. In the Khodayari model, all reactions are bi-uni or uni-uni reactions implemented by mass-action kinetics, while the Boecker and Chassagnole models use the generalized Michaelis-Menten type rate laws. Also, the relationship between the response coefficients of the original model and the linearized model highlights the differences between the models (Fig. S1). If the models were somewhat effectively similar, the scatter plots of the response coefficient of the original- and linearized model should look similar among the three models. However, the three panels show completely different trends. Thus, the three models have less similarity even when they are linearized around the steady states.

> *(4) Can the authors share their insights on what could be the underlying reasons for the bimodal distribution in Figure 1E? Even after adding random reactions, the distribution still has two modes - why is that?*

We have not yet resolved why only the Khodayari model shows the bimodal distribution of the response coefficient. However, by examining the time courses, the dynamics of the Khodayari model look like those of the excitable systems. This feature may contribute to the bimodal distribution of the response coefficient. In the future, we would like to show whether the system is indeed the excitable system and whcih reactions contribute to such dynamics.

> *(5) Considering the effects of the sparsity of the networks on the perturbation responses (from line 223 onwards), when we compare the three analyzed models, it is clear that the Khodayari et al. model is a superset of the other two models. Therefore, this model can be considered as, e*.*g*., *Chassagnole model with Nadd reactions (though not randomly added). Based on Figures 1b and S2, one can observe that the responses of the Khodayari models have stronger responses, which is exactly opposite to the authors’ conclusion that adding the reactions weakens the responses*.
>
> *The authors should comment on this*.

The sparsity of the network is defined by the ratio of the number of metabolites to the number of reactions. Note that the Khodayari model is a superset of the Boecker and Chassagnole models in terms of the number of reactions, but also in terms of the number of metabolites (Boecker does not have the pentose phosphate pathway, Chassagnole does not have the TCA cycle, and neither has oxyative phosphorylation). Thus, even if we manually add reactions to the Boecker model, for example, we cannot obtain a network that is equivalent to the Khodayari model. We added one sentence to clarify the point (Lines 254- 255).

> ***Recommendations for the authors***:
>
> *(1) Some typos: Line 57, remove ?; Line 134, correct “relaxation”*.

Thank you for pointing out. We fixed the typos.

> *(2) Lines 510-515, please rewrite/clarify, it is confusing what are you doing*.

We rewrote the sentences (Lines 529-532). We are sorry for the confusion.

> *(3) Line 522, where are the expressions above Leq and K*?*

Leq appears in the original paper of the Boecker model, but we decided not to use Leq. We apologize for not removing Leq from the present manuscript. The * in K* is the wildcard for representing the subscripts. We added a description for the role of “*”.

> *(4) Lines 525-530, based on the wording, it seems like you test first for 128 initial concentrations if the models converge back to the steady state and then you generate another set of 128 initial concentrations - is this what you are doing, or you simply use the 128 initial concentrations that have passed the test?*

We apologize for the confusion. We did the first thing. We have rewritten the sentence to make it clearer.

> *(5) Figure 3, caption, by “broken line,” did the authors mean “dashed line”?*

We meant dashed line. We changed “broken line” to “dashed line”.

https://doi.org/10.7554/eLife.98800.2.sa0

## References

Araki M, Cox RS, Makiguchi H, Ogawa T, Taniguchi T, Miyaoku K, Nakatsui M, Hara KY, Kondo A. (2015) M-path: a compass for navigating potential metabolic pathways Bioinformatics 31:905–911

Awazu A, Kaneko K. (2009) Ubiquitous “glassy” relaxation in catalytic reaction networks Physical Review E 80

Bar-Even A, Noor E, Lewis NE, Milo R. (2010) Design and analysis of synthetic carbon fixation pathways Proc Natl Acad Sci U S A 107:8889–8894

Basan M et al. (2020) A universal trade-off between growth and lag in fluctuating environments Nature 584:470–474

Biselli E, Schink SJ, Gerland U. (2020) Slower growth of Escherichia coli leads to longer survival in carbon starvation due to a decrease in the maintenance rate Molecular Systems Biology 16

Boecker S, Slaviero G, Schramm T, Szymanski W, Steuer R, Link H, Klamt S. (2021) Deciphering the physiological response of Escherichia coli under high ATP demand Mol Syst Biol 17

Calcott PH, Postgate JR (1972) On substrate-accelerated death in Klebsiella aerogenes J Gen Microbiol 70:115–122

Chakrabarti A, Miskovic L, Soh KC, Hatzimanikatis V. (2013) Towards kinetic modeling of genome-scale metabolic networks without sacrificing stoichiometric, thermodynamic and physiological constraints Biotechnol J 8:1043–1057

Chassagnole C, Noisommit-Rizzi N, Schmid JW, Mauch K, Reuss M. (2002) Dynamic modeling of the central carbon metabolism of Escherichia coli Biotechnology and bioengineering 79:53–73

Claassens NJ, Burgener S, Vögeli B, Erb TJ, Bar-Even A. (2019) A critical comparison of cellular and cell-free bioproduction systems Curr Opin Biotechnol 60:221–229

Doerr A, de Reus E, van Nies P, van der Haar M, Wei K, Kattan J, Wahl A, Danelon C. (2019) Modelling cell-free RNA and protein synthesis with minimal systems Phys Biol 16

Feinberg M. (2019) Foundations of Chemical Reaction Network Theory Cham: Springer

Ferrell JE, Ha SH (2014) Ultrasensitivity part I: Michaelian responses and zero-order ultrasensitivity Trends in biochemical sciences 39:496–503

Friedman N, Cai L, Xie XS (2006) Linking stochastic dynamics to population distribution: an analytical framework of gene expression Phys Rev Lett 97

Furusawa C, Kaneko K. (2003) Zipf’s law in gene expression Physical review letters 90

Furusawa C, Kaneko K. (2012) Adaptation to optimal cell growth through self-organized criticality Phys Rev Lett 108

Gutenkunst RN, Waterfall JJ, Casey FP, Brown KS, Myers CR, Sethna JP (2007) Universally sloppy parameter sensitivities in systems biology models PLoS Comput Biol 3:1871–1878

Haiman ZB, Zielinski DC, Koike Y, Yurkovich JT, Palsson BO (2021) MASSpy: Building, simulating, and visualizing dynamic biological models in Python using mass action kinetics PLoS Comput Biol 17

van Heerden JH, Wortel MT, Bruggeman FJ, Heijnen JJ, Bollen YJM, Planqué R, Hulshof J, O’Toole TG, Wahl SA, Teusink B. (2014) Lost in transition: start-up of glycolysis yields subpopulations of nongrowing cells Science 343

Heinrich R, Rapoport TA (1974) A linear steady-state treatment of enzymatic chains. General properties, control and effector strength Eur J Biochem 42:89–95

Himeoka Y, Gummesson B, Sørensen MA, Svenningsen SL, Mitarai N. (2022) Distinct Survival, Growth Lag, and rRNA Degradation Kinetics during Long-Term Starvation for Carbon or Phosphate mSphere 7

Himeoka Y, Kaneko K. (2017) Theory for transitions between exponential and stationary phases: universal laws for lag time Physical Review X 7

Himeoka Y, Kirkegaard JB, Mitarai N, Krishna S. (2022) Structural determinants of relaxation dynamics in chemical reaction networks bioRxiv

Himeoka Y, Mitarai N. (2022) Emergence of growth and dormancy from a kinetic model of the Escherichia coli central carbon metabolism Phys Rev Res 4

Homma T, Saltelli A. (1996) Importance measures in global sensitivity analysis of nonlinear models Reliab Eng Syst Saf 52:1–17

The MathWorks Inc. (2022) The MathWorks Inc. MATLAB version: 9.12.0 (R2022a). Natick, Massachusetts, United States: The MathWorks Inc.; 2022. https://www.mathworks.com. Natick, Massachusetts, United States: The MathWorks Inc

Kaneko K. (1981) Adiabatic elimination by the eigenfunction expansion method Progress of Theoretical Physics 66:129–142

Kaneko K, Furusawa C, Yomo T. (2015) Universal relationship in gene-expression changes for cells in steady-growth state Physical Review X 5

Kaplan Y, Reich S, Oster E, Maoz S, Levin-Reisman I, Ronin I, Gefen O, Agam O, Balaban NQ (2021) Observation of universal ageing dynamics in antibiotic persistence Nature 600:290–294

Khodayari A, Zomorrodi AR, Liao JC, Maranas CD (2014) A kinetic model of Escherichia coli core metabolism satisfying multiple sets of mutant flux data Metabolic engineering 25:50–62

Kobayashi TJ, Loutchko D, Kamimura A, Horiguchi S, Sughiyama Y. (2022) Information Geometry of Dynamics on Graphs and Hypergraphs arXiv

Kochanowski K, Volkmer B, Gerosa L, Haverkorn van Rijsewijk BR, Schmidt A, Heinemann M. (2013) Functioning of a metabolic flux sensor in Escherichia coli Proc Natl Acad Sci U S A 110:1130–1135

Kondo Y, Kaneko K. (2011) Growth states of catalytic reaction networks exhibiting energy metabolism Physical Review E 84

Kotte O, Zaugg JB, Heinemann M. (2010) Bacterial adaptation through distributed sensing of metabolic fluxes Molecular systems biology 6

Kresnowati MTAP, van Winden WA, Almering MJH, ten Pierick A, Ras C, Knijnenburg TA, Daran-Lapujade P, Pronk JT, Heijnen JJ, Daran JM (2006) When transcriptome meets metabolome: fast cellular responses of yeast to sudden relief of glucose limitation Mol Syst Biol 2

Kurisu M, Katayama R, Sakuma Y, Kawakatsu T, Walde P, Imai M. (2023) Synthesising a minimal cell with artificial metabolic pathways Commun Chem 6:1–14

Lee Y, Lafontaine Rivera JG, Liao JC (2014) Ensemble Modeling for Robustness Analysis in engineering non-native metabolic pathways Metab Eng 25:63–71

Levin-Reisman I, Gefen O, Fridman O, Ronin I, Shwa D, Sheftel H, Balaban NQ (2010) Automated imaging with ScanLag reveals previously undetectable bacterial growth phenotypes Nature Methods 7

Monk JM et al. (2017) i ML1515, a knowledgebase that computes Escherichia coli traits Nature biotechnology 35:904–908

Moriya Y, Shigemizu D, Hattori M, Tokimatsu T, Kotera M, Goto S, Kanehisa M. (2010) PathPred: an enzyme-catalyzed metabolic pathway prediction server Nucleic Acids Res 38:W138–43

Okamoto M, Hayashi K. (1983) Dynamic behavior of cyclic enzyme systems J Theor Biol 104:591–598

Okamoto M, Katsurayama A, Tsukiji M, Aso Y, Hayashi K. (1980) Dynamic behavior of enzymatic system realizing two-factor model J Theor Biol 83:1–16

Okano H, Hermsen R, Kochanowski K, Hwa T. (2019) Regulation underlying hierarchical and simultaneous utilization of carbon substrates by flux sensors in Escherichia coli Nature Microbiology 5:206–215

Olivier BG, Snoep JL (2004) Web-based kinetic modelling using JWS Online Bioinformatics 20:2143–2144

Opgenorth PH, Korman TP, Bowie JU (2014) A synthetic biochemistry molecular purge valve module that maintains redox balance Nat Commun 5

Opgenorth PH, Korman TP, Iancu L, Bowie JU (2017) A molecular rheostat maintains ATP levels to drive a synthetic biochemistry system Nat Chem Biol 13:938–942

Orth JD, Thiele I, Palsson BØ. (2010) What is flux balance analysis? Nature biotechnology 28:245–248

Paulsson J, Ehrenberg M. (2000) Random signal fluctuations can reduce random fluctuations in regulated components of chemical regulatory networks Phys Rev Lett 84:5447–5450

Pirt S. (1965) The maintenance energy of bacteria in growing cultures Proceedings of the Royal Society of London Series B Biological Sciences 163:224–231

Postgate JR, Hunter JR (1964) Accelerated death of Aerobacter aerogenes starved in the presence of growth-limiting substrates J Gen Microbiol 34:459–473

Radzikowski JL, Vedelaar S, Siegel D, Ortega ÁD, Schmidt A, Heinemann M. (2016) Bacterial persistence is an active σS stress response to metabolic flux limitation Molecular systems biology 12

Risken H., Risken H (1996) Fokker-Planck Equation The Fokker-Planck Equation: Methods of Solution and Applications Berlin, Heidelberg: Springer Berlin Heidelberg :63–95

Rizk ML, Liao JC (2009) Ensemble modeling for aromatic production in Escherichia coli PloS one 4

Rizzi M, Baltes M, Theobald U, Reuss M. (1997) In vivo analysis of metabolic dynamics in Saccharomyces cerevisiae: II Mathematical model. Biotechnol Bioeng 55:592–608

Savageau MA (1988) Introduction to S-systems and the underlying power-law formalism Math Comput Model 11:546–551

Schaechter M, Maaløe O, Kjeldgaard NO (1958) Dependency on medium and temperature of cell size and chemical composition during balanced growth of Salmonella typhimurium Microbiology 19:592–606

Schwander T, Schada von Borzyskowski L, Burgener S, Cortina NS, Erb TJ (2016) A synthetic pathway for the flxation of carbon dioxide in vitro Science 354:900–904

Scott M, Gunderson CW, Mateescu EM, Zhang Z, Hwa T. (2010) Interdependence of cell growth and gene expression: origins and consequences Science 330:1099–1102

Segler MHS, Preuss M, Waller MP (2018) Planning chemical syntheses with deep neural networks and symbolic AI Nature 555:604–610

Sekar K, Linker SM, Nguyen J, Grünhagen A, Stocker R, Sauer U. (2020) Bacterial Glycogen Provides Short-Term Benefits in Changing Environments Appl Environ Microbiol 86

Shimizu Y, Inoue A, Tomari Y, Suzuki T, Yokogawa T, Nishikawa K, Ueda T. (2001) Cell-free translation reconstituted with purified components Nat Biotechnol 19:751–755

Strange RE, Hunter JR (1966) Substrate-accelerated death’ of nitrogen-limited bacteria J Gen Microbiol 44:255–262

Tan Y, Liao JC (2012) Metabolic ensemble modeling for strain engineers Biotechnology journal 7:343–353

Taniguchi Y, Choi PJ, Li GW, Chen H, Babu M, Hearn J, Emili A, Xie XS (2010) Quantifying E coli proteome and transcriptome with single-molecule sensitivity in single cells. Science 329:533–538

Teusink B, Walsh MC, van Dam K, Westerhoff HV (1998) The danger of metabolic pathways with turbo design Trends Biochem Sci 23:162–169

Theobald U, Mailinger W, Baltes M, Rizzi M, Reuss M. (1997) In vivo analysis of metabolic dynamics in Saccharomyces cerevisiae : I Experimental observations. Biotechnol Bioeng 55:305–316

Tran LM, Rizk ML, Liao JC (2008) Ensemble modeling of metabolic networks Biophysical journal 95:5606–5617

Varma A, Palsson BO (1993) Metabolic capabilities of Escherichia coli: I Synthesis of biosynthetic precursors and cofactors. Journal of theoretical biology 165:477–502

Walther T, Novo M, Rössger K, Létisse F, Loret MO, Portais JC, François JM (2010) Control of ATP homeostasis during the respiro-fermentative transition in yeast Mol Syst Biol 6

